# Autopolyploid establishment depends on life history strategy and the mating outcomes of clonal architecture

**DOI:** 10.1101/2021.10.21.465190

**Authors:** Wendy E. Van Drunen, Jannice Friedman

## Abstract

Polyploidy is a significant component in the evolution of many taxa, particularly plant groups. However, the mechanisms promoting or preventing initial polyploid establishment in natural populations are often unclear. We develop spatially explicit agent-based simulation models to explore how perennial life history and clonal propagation influence the early stages of polyploid establishment. Our models show that polyploid establishment is unlikely among short-lived plants. Polyploids have increased establishment probability when both diploid and polyploid lifespans are long, especially when unreduced gamete production is non-zero. Further, polyploids that combine sexual and clonal reproduction can establish across a wide range of life history strategies. Polyploid genets containing many, far spreading ramets are most successful, but genets with tightly clumped ramets have similar establishment probability when pollen dispersal is local and rates of self-fertilization are high. Clonal architecture has a substantial impact on the spatial structure of the mixed cytotype population during establishment; altering patterns of mating within or between cytotypes, the mechanisms through which polyploid establishment proceeds, and the final composition of the polyploid population after successful establishment. Overall, our findings provide insight into the complex relationship between polyploidy, perenniality, and clonal reproduction, and offer testable predictions for future empirical work.

## INTRODUCTION

Polyploidy has played a key role in the evolutionary history and diversification of plants (Wood et al. 2009; Husband et al. 2013), and there is growing evidence that it is also common within many animal clades (Otto and Whitton 2000). Polyploids arise through whole-genome duplication (WGD) events resulting in more than two complete chromosome sets in an individual, which may occur through the hybridization of two species (allopolyploidy) or within a single species (autopolyploidy). Because a WGD event can result in instant reproductive isolation and drastic phenotypic change between a polyploid and its progenitors (Baack et al. 2015), polyploidy is often viewed as one of the quintessential examples of sympatric speciation. Indeed, approximately 15% and 30% of speciation events in angiosperms and ferns respectively have been attributed to polyploidization (Wood et al. 2009). Though polyploidy is prevalent throughout plants and the wider tree of life, many of the ecological and evolutionary mechanisms leading to the establishment success or failure of newly formed polyploids in natural populations are poorly understood (Husband 2000; Ramsey and Ramsey 2014; Soltis et al. 2016*b*).

The rate of polyploid formation surely exceeds the rate of successful establishment (Ramsey and Schemske 2002), because rare polyploids experience significant fitness disadvantages when they first arise in populations dominated by their lower-ploid parents (Husband 2000; Ramsey and Schemske 2002). Here, the majority of the polyploid mating opportunities are intercytotype, resulting in inviable odd-ploidy offspring and low fitness, and ultimately polyploid exclusion from the population (Levin 1975). This process of Minority Cytotype Exclusion (MCE) is expected to be a significant barrier to polyploid evolution (Thompson and Lumaret 1992; Husband 2000; Fowler and Levin 2016), and in many ways is analogous to the challenges faced by a mutant genotype invading a resident population with a lack of compatible mates, or as individuals colonize new habitat (Baack 2005; Pannell et al. 2015). In general, polyploid establishment is more probable when the effects of MCE are mitigated (Stebbins 1950), which may occur through: 1) a shift in the balance of intra- vs. intercytotype mating, 2) a reduced reliance on sexual reproduction, 3) high rates of polyploid formation, or 4) increased persistence time.

The process of polyploid establishment and MCE are challenging to study empirically (e.g., Husband 2000; Sutherland et al. 2020), but previous theoretical treatments largely confirm that polyploid establishment is facilitated by factors contributing to the four MCE-reducing processes listed above. Studies have identified important roles for self-fertilization (Levin 1975; Rodríguez 1996; Rausch and Morgan 2005), local dispersal (Baack 2005), asexuality (Nakayama et al. 2002; Yamauchi et al. 2004; Spoelhof et al. 2020), elevated unreduced gamete (UG) production (Felber and Bever 1997; Burton and Husband 2001; Husband 2004), and iteroparity (Rodríguez 1996). Most of these studies have explicitly represented plant systems and autopolyploidy (but see Fowler and Levin 2016), but even within this framework there remain significant gaps in our knowledge of the critical early period immediately following polyploid formation (Spoelhof et al. 2017).

A perennial life history strategy and asexual reproduction through clonal propagation have long been hypothesized to increase polyploid establishment success (Gustafsson 1948; Stebbins 1950; Rodríguez 1996; Chrtek et al. 2017), and both have the potential to reduce MCE via one or more mechanisms. Perenniality (coupled with iteroparity) increases the persistence of a new polyploid over several generations and the number of reproductive bouts during an individual’s lifetime, and should increase the likelihood of viable intracytotype mating via self-fertilization (Gustafsson 1948; Otto and Whitton 2000; Rice et al. 2019). Additionally, perenniality and longer polyploid lifespans may influence the probability that neopolyploid individuals originating from different WGD events will overlap in time, increasing opportunities for successful polyploid outcrossing.

Unlike perenniality, clonal reproduction directly contributes to polyploid fitness through the production of clonal offspring. Clonal propagation decreases the probability of complete polyploid exclusion, and increases the probability of intracytotype mating through geitonogamous self-fertilization between shoots within a genetic clone (i.e., a genet, comprised of numerous genetically-identical ramets; Vallejo-Marín et al. 2010). Since mating in plants generally occurs between near neighbours, the placement and arrangement of ramets can dramatically alter the sexual fitness of clonal plants by affecting pollen transport within and between genets (Handel 1985; Vandepitte et al. 2013; Van Drunen et al. 2015). The potential for geitonogamy to benefit an establishing polyploid is then inherently dependent on both population spatial structure and clonal architecture. Dense genets with closely packed ramets (i.e., phalanx architecture; Charpentier 2002; Vallejo-Marín et al. 2010) exhibit high rates of geitonogamous self-fertilization and little outcrossing. If the costs of inbreeding are low (Husband et al. 2008), a polyploid with a phalanx strategy can create its own local pocket of same-cytotype mates (Li et al. 2004; Baack 2005). In contrast, sparse genets with widely dispersed ramets (i.e., guerrilla architecture Charpentier 2002) should promote intermingling between different genets and mating via outcrossing. A guerrilla architecture and a high capacity for lateral growth could promote the spread of polyploids throughout a population (Van Drunen et al. 2015; Herben et al. 2017), and may be a better strategy if inbreeding depression is high. Thus, though either type of clonal architecture could promote polyploid establishment, the pathway to establishment could be fundamentally different between strategies. However, only a handful of polyploid establishment models include spatial or population structure (Li et al. 2004; Baack 2005; Spoelhof et al. 2020; Griswold 2021), and none have considered the effects of geitonogamy or clonal architecture.

Evaluating the influence that perenniality or clonality have on polyploid establishment is complicated by the correlations amongst these traits. At a broad scale, phylogenetic and biogeographical studies have demonstrated a positive evolutionary association between polyploidy and perenniality or clonality (Gustafsson 1948; Herben et al. 2017; Rice et al. 2019; Van Drunen and Husband 2019), but these relationships may be confounded by the fact that all clonal plants are perennial, and that the perennating organs of many plants are themselves clonal modules (e.g., tillering grasses; Klimeš et al. 1997). Thus, the relative importance of perenniality and clonality, and their potential interactions, during the initial stages of polyploid evolution are not well-defined.

Successful polyploid establishment generally requires some difference between cytotypes that conveys a benefit to the polyploids (Levin 1975), an idea corroborated by classic two-species coexistence theory (reviewed in Barabás et al. 2018). Many models incorporate this through niche shifts reducing competition between cytotypes (e.g., Rodríguez 1996), or by directly setting polyploid fitness higher than that of diploids (e.g., Baack 2005). Relatively few explicitly explore other phenotypic differences between cytotypes (but see Rausch and Morgan 2005; Chrtek et al. 2017; Griswold 2021), though in practice WGD can result in sweeping changes to gene expression (Levin 2002; Soltis et al. 2016*a*), physiology (Maherali et al. 2009; Anneberg and Segraves 2020), or morphology in new polyploids (Husband et al. 2016). WGD is expected to produce bigger cells with long cell cycles, which should slow development, delay maturity, and potentially increase lifespan (Bennett 1972; Levin 2002; Beaulieu et al. 2008; Blomme et al. 2014), resulting in polyploids that are more perennial than diploids. The immediate effects of WGD on clonal reproduction are less predictable. Newly synthesized polyploids can be more or less clonal than diploids (Van Drunen and Husband 2018*a*, 2018*b*; Walczyk and Hersch-Green 2019), and established polyploids also show a similarly variable relationship with clonality (e.g., Hroudová and Zákravský 1993; Keeler 2004; Baldwin and Husband 2013). In any case, we may hypothesize that an increase in clonal reproduction will aid polyploid establishment, while a decrease will almost certainly ensure failure for a polyploid among highly clonal diploids.

Theory is a valuable tool in exploring the elusive early steps in polyploid evolution, and for understanding how the traits implicated in these high-level patterns might causally influence key individual- and population-level processes. Here, we explore the effects of perenniality and clonal reproduction on the early stages of autopolyploid establishment using spatially explicit, agent-based simulation models incorporating the complex dynamics between reproductive strategy, perenniality, and population spatial structure. Specifically, we ask: 1) Does a perennial life history increase polyploid establishment? 2) Does clonal reproduction increase polyploid establishment? 3) How is polyploid establishment influenced by genet size and architecture? and 4) Do differences in life history or reproductive strategy between polyploids and diploids further influence these processes?

## METHODS

### MODEL OVERVIEW

Models were written in R v3.6.2 (R Development Core Team 2019), the full code is available online (https://github.com/wevandrunen/autopolyploid-establishment-lifehistory-clonalarchitecture). Simulations take place on a square population lattice measuring D x D units. The population lattice exhibits periodic boundary conditions, such that pollen, seed, or clonal ramets that disperse beyond the grid limits in one direction re-enter the population on the opposite edge (i.e., the grid is toroidal). Individuals occupy all cells in the lattice. and population size (N = D x D) is constant over each generation. Models are initialized with N – 1 diploid individuals, and a single tetraploid individual at a random location. The population then undergoes a series of biological processes over a set number of time-steps, equivalent to growing seasons or generations. For each generation, the sequence of events is: sexual and clonal reproduction, survival, and offspring recruitment (fig. 1).

**Figure 1:**
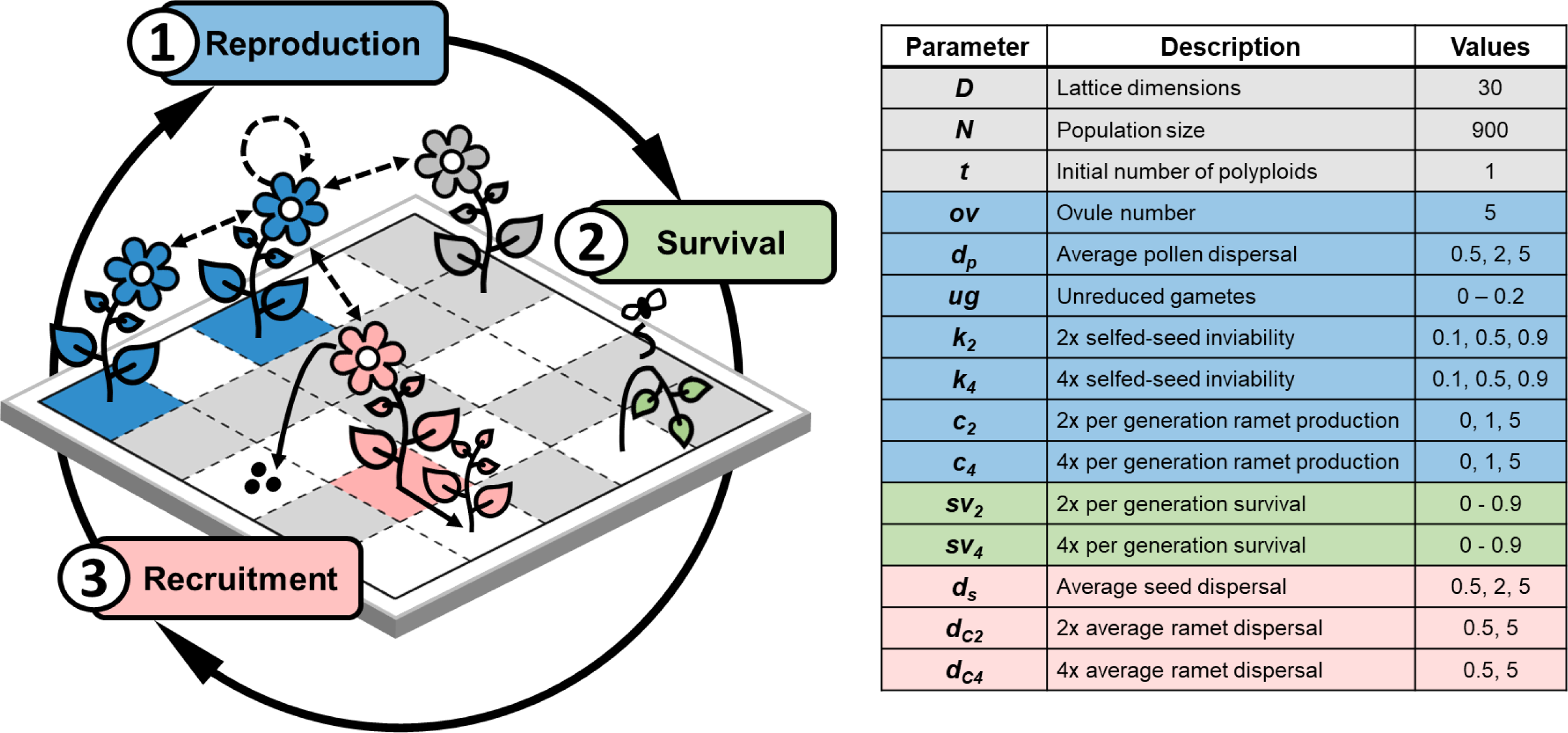
Simulation model workflow, and the adjustable parameters associated with each of the three main steps per generation. Models are initiated on a *D* x *D* population lattice containing both tetraploids (*t* individuals, 4x, coloured cells) and diploids (*N – t* individuals, 2x, grey cells), where each lattice cell can hold one individual. In Step 1, pollen is dispersed (dashed lines) within shoot, between shoots in the same genet (blue tetraploid genet), between members of the same cytotype (blue and pink tetraploids), and between cytotypes (blue tetraploid and grey diploid). Seeds are fertilized, and clonal ramets are produced. Individuals die in Step 2 according to their survival probability, leaving empty spaces in the population (white cells). In Step 3, empty cells are colonized by newly produced seeds or ramets. The surviving individuals and the new cohort of recruits then repeat this cycle, which continues for a specified number of generations. See the main text for more detail.

We make several assumptions about the equivalency of diploid vs. tetraploid individuals. Cytotypes are equal competitors and occupy the same ecological niche, and diploids and tetraploids have equal seed germination and ramet establishment probability. Finally, individual fecundity and survival are not age related; individuals reproduce sexually and asexually immediately upon recruitment into the population, have a fixed reproductive output per generation, and have a constant probability of survival to the next generation over their lifetime.

### STEP 1 – REPRODUCTION

The first step in each generation is pollen dispersal and ovule fertilization (fig. 1). All individuals are hermaphroditic and produce ov ovules per generation (fig. 1). Pollen production is unlimited and no ovule is left unfertilized, but not all fertilizations will result in viable seed (see below). Ovule fertilization proceeds by first determining the pollen density from each individual in the population at the location of the ovule source, including pollen produced by the ovule source itself. Pollen dispersal is described by Gamma distribution P(x) (eq. [1]) with shape parameter β set to 1, corresponding to leptokurtic pollen distribution (fig. S1A; figs. S1–S12 and table S1 are available online) where pollen density is highest at the pollen source (x = 0).

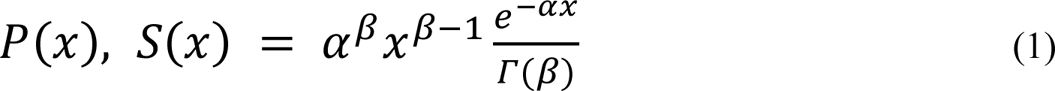

The rate parameter *α* is calculated as *β*/*d_p_*, where *d_p_* is the average pollen dispersal distance (fig. 1). Due to the toroidal shape of the population grid, pollen from an individual could arrive at a location through multiple routes, and we calculate pollen densities using the shortest distance between the pollen donor and ovule source. The selected pollen donor for each ovule is chosen by the weighted pollen probability density at the ovule location. All pollen is assumed to be equally able to fertilize an ovule: there is no pollen precedence, female mate choice, or performance differences between reduced (1n) and unreduced (2n) pollen.

The outcome of each mating event, and seed viability, is determined by the identity of the gamete sources. A mating event may result in one of 22 cross-types, that fall into four categories per cytotype: within shoot self-fertilization (autogamy), between ramet self-fertilization (geitonogamy), intercytotype outcrossing, and intracytotype outcrossing (table S1). Rates of self-fertilization vs. outcrossing are therefore emergent properties of the model, determined intrinsically through pollen donor selection. Self-fertilized seeds are inviable with probability *k_2_* or *k_4_* and viable with probability (1 - *k_2_*) or (1 – *k_4_*), for diploids and tetraploids respectively (table S1, fig. 1), which may be interpreted as early acting inbreeding depression. Diploids produce unreduced gametes (2n) with probability *ug*, and reduced gametes (1n) with probability (1 – *ug*) (table S1, fig. 1). Tetraploids make only reduced gametes (2n), and the formation of ploidy levels higher than tetraploid cannot occur. Intercytotype crosses, or intracytotype crosses involving unreduced gametes, are viable only when the offspring is tetraploid, and triploidy is assumed to be lethal (table S1). All seeds produced via intracytotype outcrossing with reduced gametes are viable. After fertilization, all viable seeds are stored in a common pool for recruitment into the next generation.

Individuals can produce a number of clonal ramets in each generation, *c_2_* for diploids, and *c_4_* for tetraploids (fig. 1). The *c_2_* and *c_4_* parameters are constant throughout a single model run, though they may differ from each other. For some scenarios, *c_2_* and/or *c_4_* are zero, meaning no ramets are produced by individuals of that cytotype and reproduction is entirely sexual. Ramets are stored in the same offspring pool as the seeds for the recruitment step.

### STEP 2 – SURVIVAL

Diploid and tetraploid individuals within the population survive between generations with probability *s_v2_* or *s_v4_* (fig. 1). For annual life histories these probabilities are zero. For perennials an average of *s_v2_* diploid and *s_v4_* tetraploid individuals survive between generations, and the remainder of the population, up to N, is supplemented by new recruits from the available offspring pool.

### STEP 3 – RECRUITMENT

Offspring are recruited onto the sites newly vacated by deceased individuals, and may be either sexually produced seedlings or clonal ramets. The recruitment process is similar to ovule fertilization, where the probability of a recruit colonizing a vacant site is calculated as the seed or ramet density at the site, dispersed from their source locations. Seed probabilities follow the Gamma function *S(x)* (eq. [1]), where β is set to 1 and the rate parameter α is *β*/*d_s_*, where *d_s_* is the average seed dispersal distance (fig. 1). In contrast, ramet dispersal *C(x)* is assumed to conform to a bell-shaped Gaussian distribution (eq. [2], variance σ^2^ = 0.15; fig. S1B) such that peak probability density occurs at the mean ramet dispersal distance from the parental source (*d_C2_* or *d_C4_*, fig. 1).

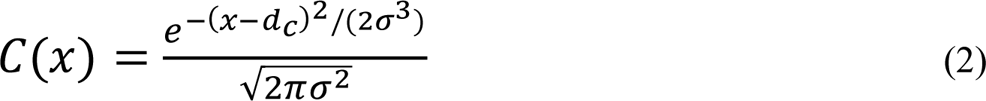

Recruitment sites are filled in random order, and offspring are removed from the common pool following successful recruitment. Since ramet probability density is formulated to be high relative to seed density at *x* ≈ *d_c_* from a source plant (fig. S1), clonal recruitment will occur much more frequently at these distances than seed colonization, mirroring a biologically realistic scenario where the competitive and establishment ability of ramets is higher than that of seedlings (Vallejo-Marín et al. 2010). Centering ramet colonization at *d_c_* results in higher ramet density at a “spacer distance” away from the primary shoot and imposes strong spatial structure on clonal genets (e.g., Winkler and Stöcklin 2002). After recruitment, seedlings are assigned new unique genet IDs, while ramets retain the ID of their parent. Data on the survivors and new cohort of individuals is passed to the next generation, and the reproduction step begins again.

### PARAMETERIZATION

There are 15 adjustable parameters in the model (fig. 1). Except where noted, simulations take place on a square lattice with dimension *D* = 30 and a population of *N* = 900, which remains constant across generations. Simulations begin with only one polyploid individual (*t* = 1). The number of ovules produced per individual (*ov*) is five for both diploids and tetraploids, resulting in 4500 seeds per generation for a population of *N* = 900. Under the most restrictive circumstances for sexual reproduction (i.e., high selfed-seed inviability, limited pollen dispersal, no unreduced gamete production, and similar cytotype frequencies), trials showed that at least 2000 viable seeds were produced per generation; enough to minimize the possibility of anomalous long-distance dispersal events. Diploid individuals may produce unreduced gametes (UGs) at frequency *ug*, and reduced gametes at frequency 1 – *ug* (table S1).

Two parameters affect the realized amount of self-fertilization: pollen dispersal distance (*d_p_*) alters the probability of self-pollination (autogamous or geitonogamous), and selfed-seed inviability (*k*) alters the probability that self-fertilized offspring are included in the pool of viable recruits (fig. 1, table S1). We focus on these two parameters because they are expected to be influential on sexual fitness in clonal plants (Eckert 2000; Van Drunen et al. 2015), where dispersal distance determines pollen kernel density (fig. S1), and regulates whether pollen is likely to remain within a ramet, encounter other ramets in the same genet, or encounter ramets from different genets. We test the effects of nine combinations of *d_p_* and *k* (3 x 3) across different model scenarios, hereafter referred to as the core dispersal-inviability parameter set. The three core value for average pollen dispersal distance (*d_p_*) were 0.5 grid units (pollen rarely exits the source cell), 2 (local dispersal), and 5 (far dispersal) (figs. 1, S1). For simplicity, average seed dispersal distances were assumed to mirror those of pollen (*d_p_* = *d_s_*), and are equal between cytotypes for all simulations. The probability of selfed-seed inviability for diploids (*k_2_*) and tetraploids (*k_4_*) took the values 0.1, 0.5, or 0.9 (fig. 1) and were equal in most scenarios, where larger values indicate stronger inviability or inbreeding depression.

The remaining six parameters differentiate the life history and reproductive strategies of diploid or tetraploid individuals: per generation survival (*s_v2_* and *s_v4_*), clonal ramet production (*c_2_* and *c_4_*), and average ramet dispersal distance (*d_C2_* and *d_C4_*) (fig. 1). These parameters are described in more detail in the following sections.

### LIFE HISTORY STRATEGY SCENARIOS

We first examine the effect of perenniality on polyploid establishment in the absence of clonal reproduction (*c_2_* = *c_4_* = 0). Diploid and tetraploid individuals may have the same probability of survival between generations (*s_v2_* = *s_v4_*), or survival probabilities may differ (*s_v2_* ≠ *s_v4_*). For equal strategies, *s_v2_* and *s_v4_* ranged from 0 (a short-lived annual strategy with zero survival probability between generations) to 0.9 (a long-lived perennial strategy with a 90% survival probability) in increments of 0.1 (fig. 1). For the core dispersal-inviability parameter set, there were 90 unique scenarios where diploids and polyploids have equal life-histories.

Because genome duplication is expected to slow growth rates and increase lifespans, we only consider tetraploid survival probability exceeding that of diploids, defining a “Tetraploid Survival Advantage” as *s_v4_* - *s_v2_*. Diploid survival probability per generation ranged from 0 to 0.8, while tetraploid survival could take any larger value (e.g., for *s_v2_* = 0, *s_v4_* can be 0.1 – 0.9). This results in 45 pairs of *s_v2_* and *s_v4_*, and a total of 405 parameter combinations.

We ran several additional simulations to determine how alternate model formulations could influence the effect of life history on tetraploid establishment. For these models, we consider:

1. *Low population density* – Imposing a fully occupied grid may misrepresent the population structure of some plants; more closely resembling a population of long-lived trees than patchy herbaceous species. Less local competition for recruitment sites may increase polyploid establishment potential. We reran the full set of 495 life history simulations on a larger population lattice of *D* = 40 units with population size constant at *N* = 900 randomly distributed individuals, for a ∼45% decrease in plant density.
2. *Unequal selfed-seed inviability* – Because inbreeding depression is expected to be weaker in polyploids (Husband and Schemske 1997; Husband et al. 2008), we tested scenarios where the probability of selfed-seed inviability for diploids was *k_2_* = 0.9, and for tetraploids *k_4_* took lower values in increments of 0.2 (minimum 0.1). For these 4 *k_2_*-*k_4_* pairs, we reran the equal and unequal life history scenarios described above with pollen-seed dispersal *d_p_* = *d_s_* = 0.5 only (220 total parameter combinations).
3. *Non-zero UG production* – We consider values of unreduced gamete production (*ug*) by diploids of 0.01, 0.05, 0.1, 0.15, and 0.2 (fig. 1). Simulations for equal and unequal life histories were rerun for a subset of the core dispersal-inviability parameters (*d_p_* = *d_s_* = 0.5, and *k_2_ = k_4_* = 0.1), for 275 total parameter combinations.

### CLONAL REPRODUCTION SCENARIOS

Here, we allow diploid and tetraploid ramet production to be non-zero; reproduction occurs both sexually and clonally. In the equal clonality scenario, both diploids and tetraploids are perennial (*s_v2_* > 0, *s_v4_* > 0), and produce an equal number of clonal ramets per generation (*c_2_* = *c_4_* > 0) at some average distance away from the parent plant (*d_c2_* = *d_c4_* > 0). For both ramet production and dispersal we consider two values: small vs. large genets (*c_2_* = *c_4_* = 1 or 5 ramets produced per generation) and clumped vs. dispersed clonal architecture (*d_c2_* = *d_c4_* = 0.5 or 5 mean grid units). All four combinations of clonal parameters were examined, hereafter referred to as clonal strategies. For each strategy we include equal and unequal life history strategies, resulting in 1620 parameter combinations over the core dispersal-inviability set.

We vary ramet production and clonal architecture separately to determine how differences in clonality between cytotypes alters polyploid establishment, concentrating on equal life history strategies only. Keeping ramet dispersal distance (i.e., clonal architecture) equal between cytotypes (*d_c2_* = d_c4_ = 0.5 or 5), there are six combinations of *c_2_* and *c_4_*, including *c_2_* = 0 or *c_4_* = 0, for a total of 108 scenarios for the core dispersal-inviability parameter set. Similarly, keeping ramet production equal between cytotypes (*c_2_* = *c_4_* = 1 or 5) while varying ramet dispersal results in 18 additional combinations.

To test alternate model formulations for the clonal scenarios, we also consider:

1. *Exponential ramet dispersal* – Our model implicitly includes a competitive advantage to clonal ramets over seedling recruits by assuming a normal Gaussian distribution for ramet dispersal probability (eq. [2]; fig. S1B). We compared our main results with a model where ramet dispersal follows an exponential density function (eq. [1]; fig. S1A), so that the recruitment probability at a site *x* grid units away from the parent plant is the same for both sexual and clonal offspring. We reran the full set of equal clonality scenarios (1620 parameter combinations).
2. *Unequal selfed-seed inviability* – We set the probability of selfed-seed inviability for diploids at *k_2_* = 0.9, while for tetraploids *k_4_* took lower values in increments of 0.2 (minimum 0.1). For these 4 *k_2_*-*k_4_* pairs, we reran the equal clonality and equal life history scenarios described above, for ramet dispersal with either a Gaussian (432 parameter combinations) or exponential probability function (432 parameter combinations).

### MODEL OUTPUT

Simulations were replicated 20 times for each of the 5175 unique sets of parameters we investigated, for a total of 103 500 model runs. A replicate could result in three possible outcomes: exclusion of diploids (fixation of tetraploids), exclusion of tetraploids (fixation of diploids), or a stable equilibrium that includes both cytotypes. Exclusion of diploids occurs once tetraploids have replaced all diploids—because tetraploids cannot produce double-reduced gametes, there is no way for diploids to re-enter the population. Complete exclusion of tetraploids can only occur when UG production is zero; otherwise a low equilibrium frequency of tetraploids is expected as new tetraploid individuals are occasionally formed via UGs. In this case, the stability of cytotype coexistence was assessed by examining average tetraploid frequency over the last 100 generations of the simulation, and deviations of less than 10 individuals per generation over this period were deemed stable. One of the three final states was generally reached within 100 generations, but we continued simulations for a maximum of 500 generations when unreduced gamete (UG) production was non-zero, to capture rare or delayed events. To determine the probability of polyploid fixation for a particular parameter set, we calculated the proportion of replicates that end in each state, and average the number of generations needed to reach them.

For each simulation run, we tracked several demographic and reproductive metrics per generation, including diploid and tetraploid survivorship, number and average size of diploid and tetraploid individuals and genets, average local neighbourhood composition (defined as an extended Moore neighbourhood 2 grid units in each dimension; total 24 surrounding cells not including the focal cell), local neighbourhood vacancy (total 25 cells including the focal cell), the total number of offspring of each sexual cross-type produced (table S1), the number of clonal ramets produced, as well as the number of each offspring type successfully recruited into the population.

## RESULTS

### LIFE HISTORY

In models with equal survivorship (*s_v2_* = *s_v4_*) between cytotypes and no clonal reproduction, there was no successful establishment of tetraploids, unless other conditions were also met (e.g., *ug* ≠ 0). Under unequal survivorship (*s_v2_* ≠ *s_v4_*), tetraploid establishment only occurred under short pollen and seed dispersal distances (*d_p_* = *d_s_* = 0.5, high autogamy). Establishment probability increased when both diploids and tetraploids had high survival, and when tetraploids had significant survival advantages over diploids (fig. 2A). When selfed-seed inviability was high (*k_2_* = *k_4_* = 0.9), tetraploids needed a large survivorship advantage and a survival probability of at least 0.8 to establish, regardless of diploid survival (fig. 2A). When selfed-seed inviability was low (*k_2_* = *k_4_* = 0.1), tetraploids needed a minimum survival advantage of 0.4 when among annual diploids (*s_v2_* = 0), but only 0.2 when diploids had higher survival probabilities (fig. 2A). Time to tetraploid fixation was more rapid with larger tetraploid survival advantages, but was not affected by diploid survival probability; whereas tetraploid exclusion was significantly slower when tetraploid survival advantage was large and when diploid survival was high (fig. S2).

**Figure 2:**
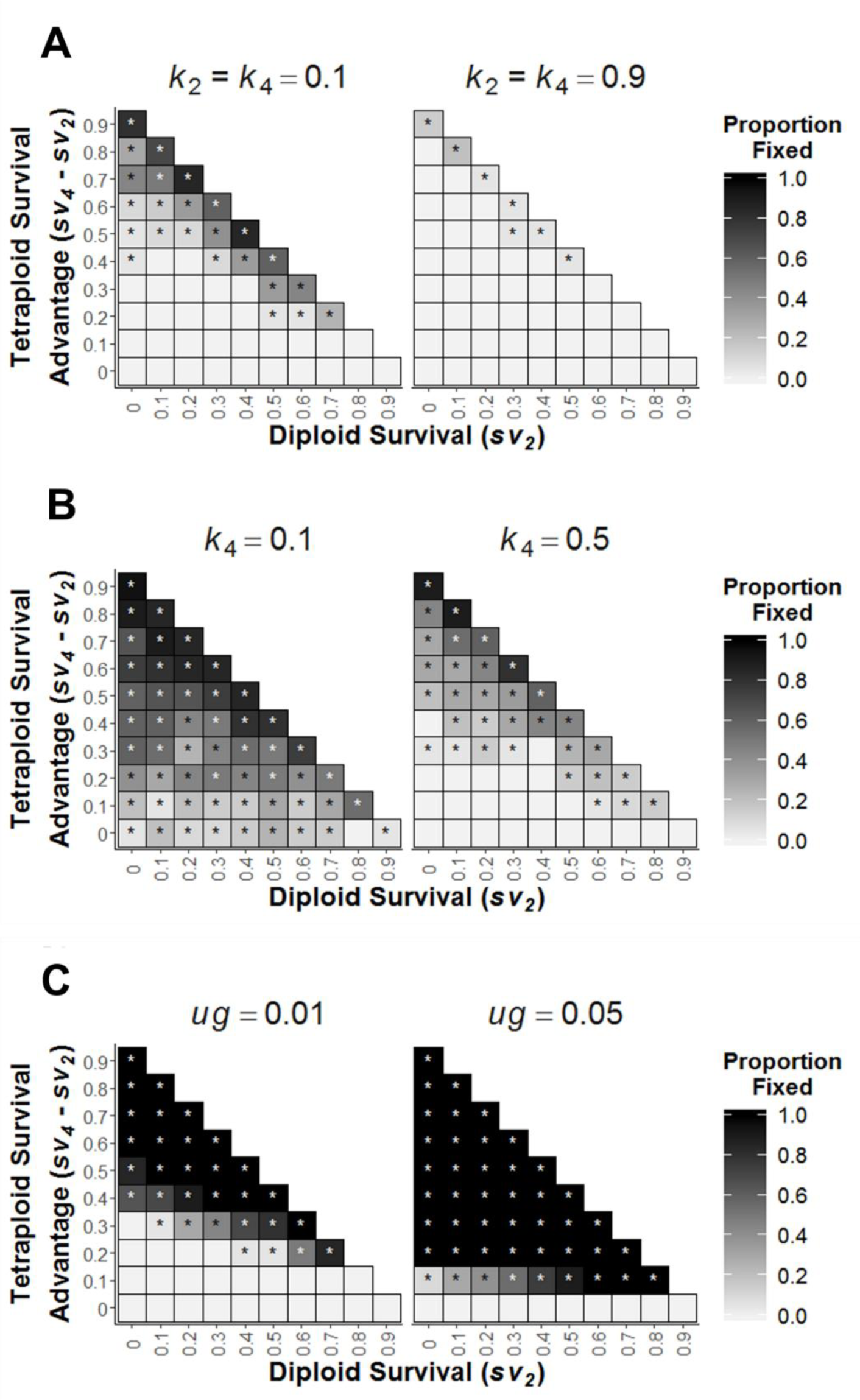
Effect of life history strategy on polyploid establishment for a range of diploid and tetraploid survival probabilities (**s_v2_*, *s_v4_**). Establishment success is measured as the proportion of simulation replicates in which tetraploids spread to fixation, and exclude diploids from the population. Panel A shows establishment under equal selfed seed inviability (*k_2_* = *k_4_*), while in Panel B tetraploid selfed-seed inviability is lower than diploids (*k_2_* = 0.9, *k_4_* value above panels). There is no unreduced gamete production in Panels A and B (*ug* = 0). In Panel C, unreduced gamete production is non-zero, and selfed-seed inviability is *k_2_* = *k_4_ =* 0.1. Asterisks denote parameter combinations where at least one simulation replicate resulted in tetraploid fixation. For all models *d_p_* = *d_s_* = 0.5.

Lower population densities expanded the parameter space that allowed successful tetraploid establishment (fig. S3A), but tetraploid fixation or exclusion times were longer (figs. S3B, C). Reducing the inviability of selfed-seed in tetraploids relative to diploids significantly increased establishment probability (fig. 2B), even when cytotype survival was equal. Times to exclusion were similar to the equal self-seed inviability simulations, but fixation times were significantly faster (figs. S4A, B).

The rate of unreduced gamete (UG) production was positively associated with tetraploid fixation probability, with the critical value being between *ug* = 0.15 and *ug* = 0.2 when cytotype life history strategies were equal and pollen-seed dispersal distance was moderate to far (*d_p_* = *d_s_* = 2 or 5), and this threshold was reduced to *ug* = 0.1 when *d_p_* = *d_s_* = 0.5 and selfed-seed inviability was low (*k_2_* = *k_4_* = 0.1; data not shown). When cytotypes varied in life history strategy, tetraploids were always successful when *ug* ≥ 0.1 (fig. S5A), unless diploid and tetraploid survival were both very high (*s_v2_* = *s_v4_* = 0.9), where 500 generations was just shy of enough time for complete fixation (fig. S6A). When *ug* = 0.05, a tetraploid survival advantage of 0.2 resulted in tetraploid establishment across all diploid survival probabilities (fig. 2C), and when tetraploid advantage was 0.1 establishment success was positively correlated with diploid survival; low diploid survival resulted in tetraploid establishment only rarely, intermediate survivals (*s_v2_* = 0.3-0.4) showed a trend towards eventual tetraploid establishment (fig. S6A), and *s_v2_* ≥ 0.5 always ended in tetraploid fixation in less than 500 generations (fig. 2C). Further reducing UG production to *ug* = 0.01 revealed a similar pattern (fig. 2C, Figure 6A). For simulations that did not proceed towards tetraploid fixation, both cytotypes coexisted with low tetraploid frequencies of ∼10 tetraploids per generation (1% of the population) for *ug* = 0.05, and ∼2 (0.2%) when *ug* = 0.01 (figs. S6A, B). In general, mean time to tetraploid fixation was slower for more perennial populations, and when tetraploid survival advantage was small, but these relationships were dependent on *ug* value (fig. S5B).

### CLONAL REPRODUCTION

#### Clonality and Life History Strategy

Tetraploid establishment probability was higher when individuals were able to reproduce both sexually and clonally (fig. 3), but the effects of clonality were strongly dependent on genet size and architecture (small vs. large genets, clumped vs. dispersed ramets). As in the nonclonal scenarios, establishment increased with both diploid survival and tetraploid survival advantage, but still no establishment occurred when cytotypes had equal life histories (figs. 3, S7). Large dispersed clonal strategies had the highest establishment probabilities across parameter space (fig. 3). Differences between clonal strategies were weaker when pollen and seed dispersal distances were short (*d_p_* = *d_s_* = 0.5, high autogamy, low geitonogamy), but when pollen and seed dispersal was far (*d_p_* = *d_s_* = 5, low autogamy and high geitonogamy) small and clumped genets were substantially less successful than large and dispersed genets (fig. 3). Selfed-seed inviability did not influence clonal tetraploid establishment probability (fig. S7), nor did lowering selfed-seed inviability for tetraploids relative to diploids (fig. S8). Mean time to tetraploid fixation was faster for all clonal strategies compared to the nonclonal, but there were no consistent differences in exclusion time (fig. S9). The effects of life history strategy on establishment under the alternate exponential formulation for ramet dispersal were nearly identical to the nonclonal scenarios, but with faster time to fixation (fig. S10). Unlike under Gaussian ramet dispersal, reducing tetraploid selfed-seed inviability compared to diploids increased establishment probability (fig. S8).

**Figure 3:**
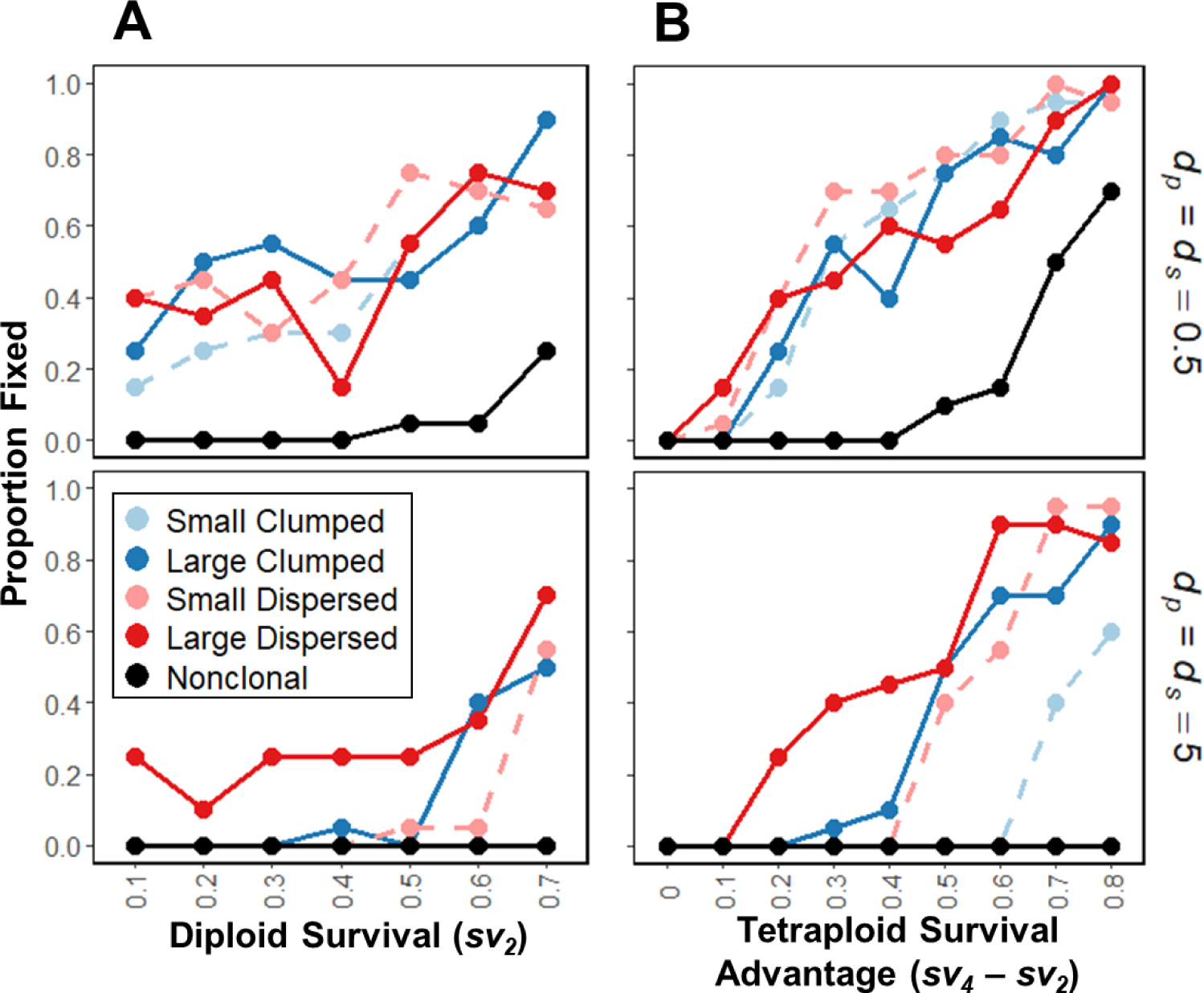
Influence of interactions between clonal strategy and life history on polyploid establishment, when cytotypes have identical clonal architectures (*Clumped d_C2_* = *d_C4_* = 0.5, blue; *Dispersed d_C2_* = *d_c4_ =* 5, red), and equal ramet production (genets are *Small c_2_* = *c_4_* = 1, light dashed lines; or *Large c_2_* = *c_4_* = 5, dark solid lines). In Panel A diploid survival probability (**s_v2_**) takes the values on the x-axis and tetraploid survival (**s_v4_**) is 0.2 higher. In panel B diploid survival (**s_v2_**) is held at 0.1, and tetraploids have a survival advantage (**s_v4_* - *s_v2_**) according to the x-axis. Average pollen (*d_p_*) and seed (*d_s_*) dispersal distances vary between rows in each panel. For all models, selfed-seed inviability is equal between cytotypes (*k_2_* = *k_4_* = 0.1), and there is no unreduced gamete production (*ug* = 0).

#### Divergent Clonal Strategies

Tetraploids with fewer ramets than diploids had no establishment success (fig. 4). When tetraploid ramet production was higher, tetraploid establishment depended on clonal architecture, pollen and seed dispersal distance, and life history strategy (fig. 4). Increasing pollen and seed dispersal distances decreased establishment probability, particularly for clumped genets (fig. 4). Tetraploids with greater survivorship tended to have lower establishment success when the difference in genet size between cytotypes was large (*c_2_* = 0, *c_4_* = 5; fig. 4), but establishment was positively correlated with survival when differences were small (*c_2_* = 0, *c_4_* = 1) and genets were dispersed (fig. 4). Selfed-seed inviability did not affect tetraploid establishment (results not shown).

**Figure 4:**
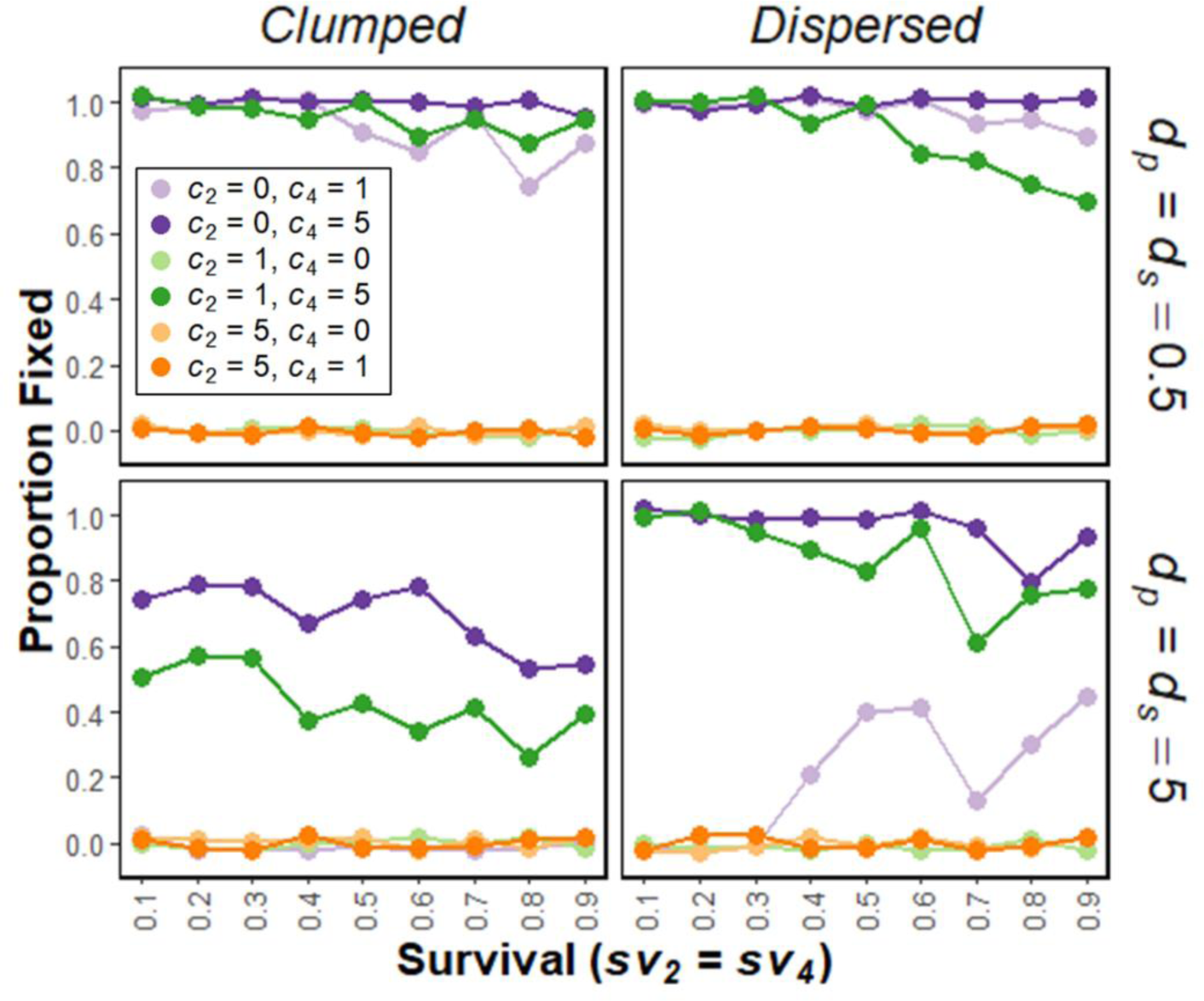
Polyploid establishment probability when diploids and tetraploids have unequal ramet production (i.e., unequal genet size, *c_2_* ≠ *c_4_*). In comparison to diploids, tetraploids can produce a few more ramets (*c_2_* = 0, *c_4_* = 1; light purple lines), many more ramets (*c_2_* = 0 or 1, *c_4_* = 5; dark purple and dark green lines), fewer ramets (*c_2_* = 5, *c_4_* = 1; dark orange lines), or no ramets (*c_2_* = 1 or 5, *c_4_* = 0; light green and light orange lines). Clonal architecture is the same for both cytotypes, with the *Clumped* strategy in the left column (*d_C2_* = *d_C4_* = 0.5), and the *Dispersed* strategy in the right column (*d_C2_* = *d_C4_* = 5). Average pollen and seed dispersal distances (*d_p_* = *d_s_*) varies between rows. Life history strategies are equal between cytotypes (**s_v2_** = **s_v4_**), as shown on the x-axis. For all models, selfed-seed inviability is equal between cytotypes (*k_2_* = *k_4_* = 0.1), and there is no unreduced gamete production (*ug* = 0). Points are slightly jittered vertically.

Tetraploids with clumped clonal architecture could not establish among dispersed diploids, but dispersed tetraploid genets generally had high establishment success among clumped diploids (fig. S11). In the latter scenario, tetraploids with large genets were little affected by pollen and seed dispersal distance, and establishment decreased with survival (fig. S11). Conversely, for small genets establishment success decreased with increasing pollen and seed dispersal distance (fig. S11). Here, increasing survival lowered establishment probability when pollen and seed dispersal was restricted (*d_p_* = *d_s_* = 0.5), but enhanced it when pollen and seed dispersal was far (*d_p_* = *d_s_* = 2 or 5, fig. S11). Selfed-seed inviability did not affect tetraploid establishment when clonal architecture differed between cytotypes (results not shown).

#### The Mating and Demographic Consequences of Clonal Reproduction

To illustrate the effects of different clonal strategies on patterns of tetraploid establishment, we focus on the set of models where *s_v2_* = 0.5, *s_v4_* = 0.9, *d_p_* = *d_s_* = 0.5 or 5, *k_2_* = *k_4_* = 0.1, and *ug* = 0, as this combination led to successful establishment in the majority of simulations. Diploids and tetraploids have equal clonal strategies (*c_2_* = *c_4_*, *d_c2_* = *d_C4_*).

Reproductive and clonal strategies substantially influenced the spatial structure of tetraploid populations, even when overall establishment probabilities were similar (fig. 5A). In the nonclonal scenarios, tetraploids spread into the diploid population in a tight circle, and at 30 generations only occupied 3-5% of the population (fig. 5B). In comparison, clonal tetraploids established quicker, with clumped architectures expanded from localized points, and dispersed architectures quickly establishing across the entire population lattice (fig. 5B). Low ramet production promoted the establishment of many distinct tetraploid genets, but populations with high ramet production were often dominated by a few very large genets (fig. 5B).

**Figure 5:**
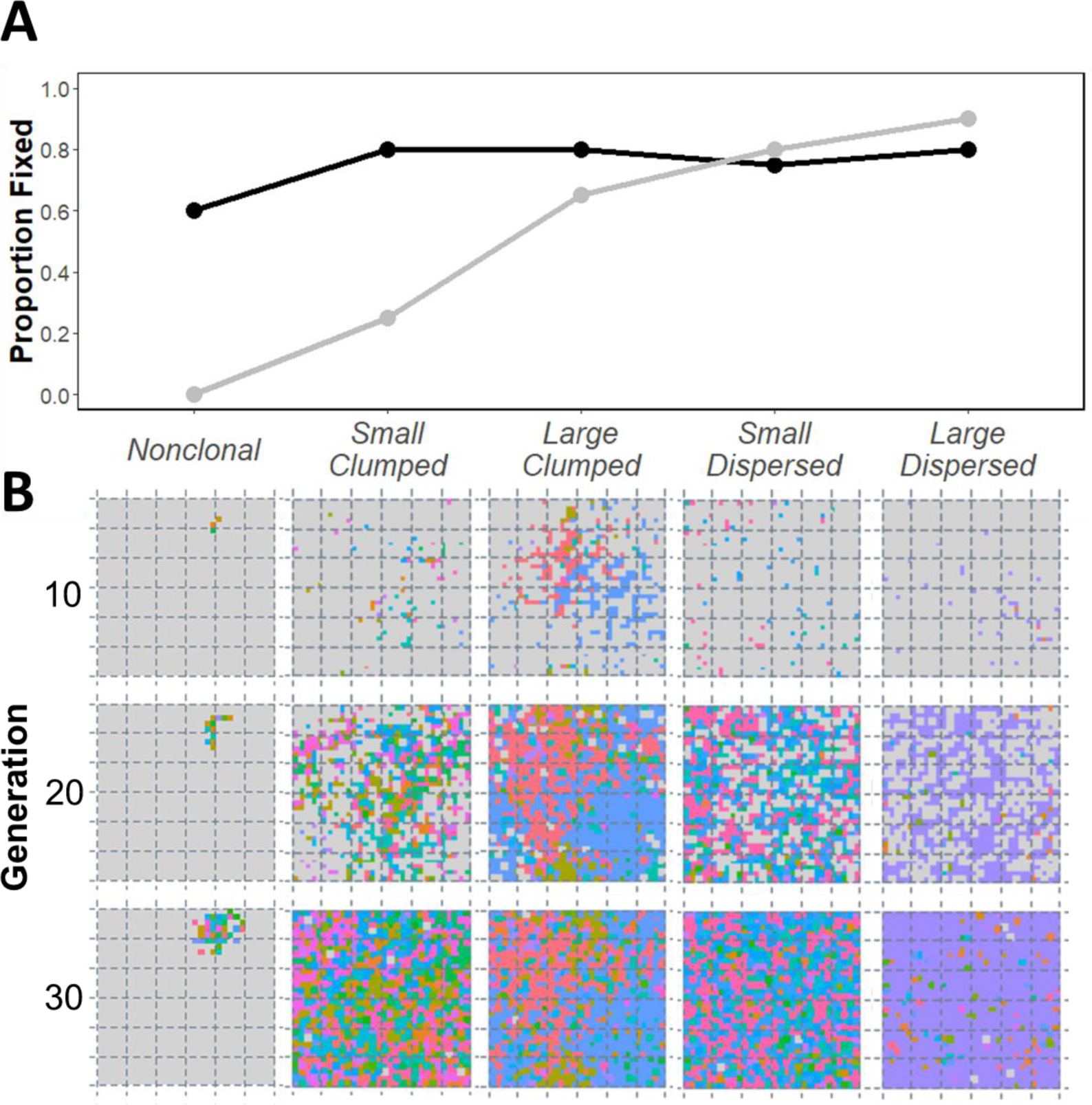
Contrasting the spread and establishment probability of nonclonal tetraploids and each clonal strategy. Panel A shows the proportion of simulations with successful polyploid establishment between nonclonal and clonal strategies (black *d_p_* = *d_s_* = 0.5, grey *d_p_* = *d_s_* = 5). Cytotypes have identical architecture (*Clumped d_C2_* = *d_C4_* = 0.5; *Dispersed d_C2_* = *d_c4_ =* 5), and equal ramet production (genets are *Small c_2_* = *c_4_* = 1; or *Large c_2_* = *c_4_* = 5). In Panel B, tetraploids (coloured cells) expand into the 2-dimensional population of diploids (grey cells), for *d_p_* = 0.5 (black line in A). Each colour represents a different tetraploid genet. For all models, **s_v2_** = 0.5, **s_v4_** = 0.9, selfed-seed inviability is equal between cytotypes (*k_2_* = *k_4_* = 0.1), and there is no unreduced gamete production (*ug* = 0).

**Figure 6:**
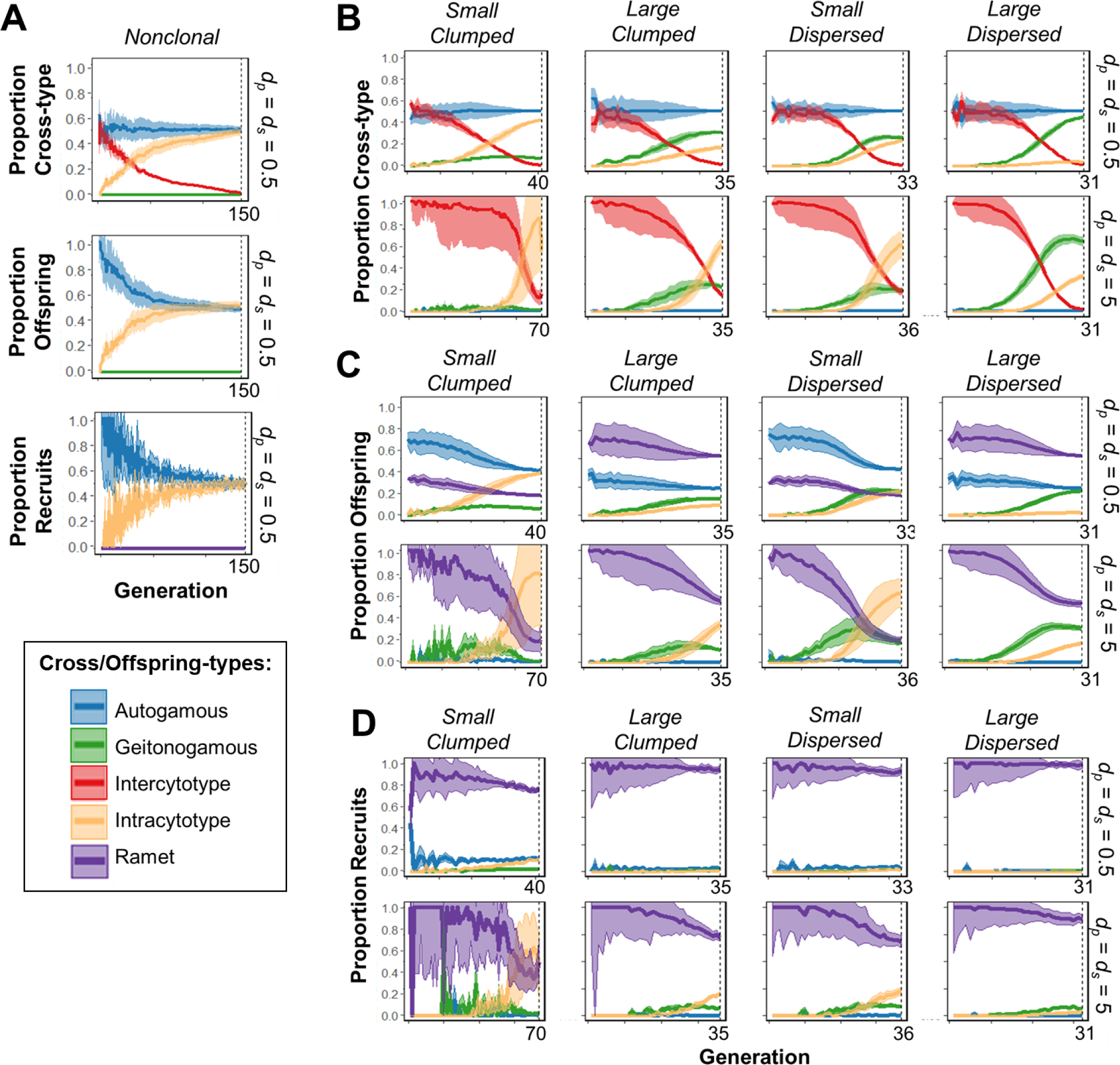
Mating outcomes and offspring recruitment during polyploid establishment, averaged (± SE) across simulation replicates per generation. Tetraploid mating events in Panel A (nonclonal) and Panel B (clonal) can result in four cross-types: autogamous self-fertilization (blue, cross-types #16-17 in Table S1), geitonogamous self-fertilization (green, #18-19), intercytotype mating (red, #20), or intracytotype mating (yellow, #22). The recruitment pool in Panel A (nonclonal) and Panel C (clonal) contains any viable sexual offspring and clonal ramets (purple) produced, which may then be recruited into the population as shown in Panel A (nonclonal) and Panel D (clonal). In Panel A all scenarios have *d_p_* = 0.5, as no establishment occurred when *d_p_* = 5. In Panels B - D, clonal strategy varies across columns, and average pollen and seed dispersal distance (*d_p_* = *d_s_*) varies between rows. Cytotypes have identical architecture (*Clumped d_C2_* = *d_C4_* = 0.5; *Dispersed d_C2_* = *d_c4_ =* 5), and equal ramet production (genets are *Small c_2_* = *c_4_* = 1; or *Large c_2_* = *c_4_* = 5). For all models, **s_v2_** = 0.5, **s_v4_** = 0.9, selfed-seed inviability is equal between cytotypes (*k_2_* = *k_4_* = 0.1), and there is no unreduced gamete production (*ug* = 0). Generations are truncated at the fastest tetraploid fixation time per parameter combination, indicated on the bottom right of each panel.

Structural differences between clonal strategies affected patterns of within- vs. between- cytotype mating during the establishment process, and the timing and location of recruitment of sexual vs. clonal offspring. For nonclonal tetraploids, when simulations began most ovules were involved in self-fertilized autogamous or inviable intercytotype matings, and nearly all viable offspring and recruits were autogamously produced (fig. 6A). As the simulations progressed and tetraploids increased in number, intracytotype matings and recruits became more common (fig. 6A).

All clonal tetraploids experienced early increases in geitonogamous mating (fig. 6B) and geitonogamous seed production (fig. 6C). When pollen and seed dispersal were restricted (*d_p_* = *d_s_* = 0.5) autogamy was prevalent, though geitonogamy remained high for large and dispersed strategies until tetraploid fixation, whereas outcrossed mating was more frequent in small and clumped genets (fig. 6B). In contrast, when pollen and seed dispersal distances were far (*d_p_* = *d_s_* = 5) geitonogamous mating was dominant early during establishment, but intercytotype mating overtook geitonogamy before fixation for all but the large dispersed clonal strategy (fig. 6B).

Sexual offspring were rarely recruited in the clonal scenarios, even when they were more abundant than ramets (figs. 6C, D). Seedling recruitment frequencies mirrored those of the viable seedlings available (fig. 6C), and was more common late during the establishment process, under far pollen and seed dispersal distances (*d_p_* = *d_s_* = 5), and for small and clumped clonal architectures (fig. 6D).

Under the alternate exponential formulation for ramet dispersal, tetraploid establishment was nearly identical to the nonclonal scenario; sexual offspring recruited were mainly the result of autogamy, and geitonogamy played little to no role (fig. S12). However, clonal ramet recruitment spiked early in establishment, particularly for large and dispersed strategies, but then remained at a low frequency until tetraploid fixation (fig. S12).

## DISCUSSION

Our results indicate that a perennial life history strategy and clonal reproduction can increase polyploid establishment potential. However, we find that this is only the case when polyploids have some advantage over diploids (e.g., higher per generation survival), or when unreduced gamete production is high, suggesting that the ability to perennate or to produce clonal ramets may not be enough to overcome Minority Cytotype Exclusion (MCE) in some circumstances. By investigating how various clonal and life history strategies affects the spatial structure, demography, and mating patterns within a mixed-ploidy population, we identify mechanisms that can influence the spread of polyploids into a diploid population. Our model is the first to incorporate the spatial attributes of clonal architecture and within-genet geitonogamous self-fertilization, and we find that clonal strategy has a significant effect on the polyploid establishment process, and major consequences for polyploid population structure.

### PERENNIALITY PROMOTES POLYPLOIDY

Perennial polyploids should experience weaker Minority Cytotype Exclusion (MCE) pressure than annual polyploids because lengthening lifespan increases the chances of reproductive success via self-fertilization, and reduces stochastic loss of all polyploids from the population. Our model supports these hypotheses and provides mechanistic links behind the broad-scale associations between polyploids and perenniality (Rice et al. 2019; Van Drunen and Husband 2019), which highlight the importance of spatial structure, temporal dynamics, and chance during the first few generations after a neopolyploid arises.

We find that polyploids are more likely to succeed among longer-lived diploid species, especially if perennial diploids give rise to polyploids with slightly longer lifespans. Notably, the polyploid survival advantage needed for consistently successful establishment decreased with increasing diploid perenniality (fig. 2A). This pattern was driven by the demographic consequences of 1) competition for space in the population, and 2) drawn-out polyploid persistence when polyploids, or both cytotypes, were long-lived. When diploids are short-lived, recruitment sites open up close to the rapidly-cycling diploids, which precludes the initial recruitment of polyploid offspring and increases the likelihood of stochastic loss of all polyploids unless they have very high relative survivorship. In contrast, when diploids and polyploids are both long-lived there is significantly slower population turnover, longer polyploid persistence, and more potential for polyploid offspring recruitment. However, when population density is low the conditions under which polyploids can establish are greater (fig. S3). This suggests that in less competitive environments, polyploid success may be less dependent on the life history strategy of diploids, in line with previous models showing that ecological niche (Fowler and Levin 1984; Rodríguez 1996) and physical (Griswold 2021) segregation can facilitate polyploid establishment, even when individuals have an annual life history strategy.

Interactions between the spatial and demographic effects of divergent life history strategies in mixed-cytotype populations offers a reason why polyploidy is more common in perennial species – the ≥40% increase in survivorship needed to establish among short-lived diploids in our model (fig. 2A) may be an unfeasibly large shift in neopolyploids, whereas a survival probability increase of 20% among long-lived diploids may be more tenable. There are numerous examples of naturally occurring polyploids that exhibit a life history shift compared to their diploid relatives. For example, in the mixed-ploidy species *Centaurea stoebe* tetraploids are iteroparous short-lived perennials while diploids are predominantly semelparous annuals (Mráz et al. 2012), and tetraploid cultivars of *Nasturtium officinale* perennate more easily than diploid lines (Manton 1935). But few studies have addressed life history changes in perenniality in neopolyploids (but see Müntzing 1936), limiting our understanding of the strength of the immediate effects of WGD and phenotypic differences between diploids and polyploids, and the consequences for polyploid establishment in natural populations. Indeed, here we demonstrate that autopolyploids are unable to establish among diploids with equal life history strategies unless they have additional advantages (e.g., significantly lower inbreeding depression), a result corroborated by previous models (Rodríguez 1996; Chrtek et al. 2017; Spoelhof et al. 2020).

Polyploid persistence time was prolonged when diploids and polyploids were highly perennial, considerably slowing model dynamics by decreasing polyploid death and recruitment rates (see also Chrtek et al. 2017; Spoelhof et al. 2020). Surveys in natural populations have found that cytotype frequencies in annual species can fluctuate wildly between growing seasons, sometimes leading to the complete loss of diploids or polyploids within a population (Čertner et al. 2017), whereas cytotype coexistence in perennial species may be maintained over many generations (Lumaret et al. 1987; Keeler 2004; Kao and Parker 2010). Thus, perenniality has been invoked as a non-adaptive factor promoting cytotype coexistence in nature (Duchoslav et al. 2010; Hanzl et al. 2014; Hanušová et al. 2019), though it is often unknown whether mixed-ploidy populations are in stable states, or represent only short snapshots of ongoing competition and exclusion (Kolář et al. 2017). Our results suggest that perenniality may contribute to an illusion of stable coexistence for a much longer timeframe than field studies typically encompass (e.g., (e.g., Čertner et al. 2017; but see Buggs and Pannell 2006). For some perennial scenarios both cytotypes coexisted for more than 300 generations while moving steadily towards polyploid fixation or exclusion (fig. S2), whereas annual polyploids were typically excluded in 1 or 2 generations. Either end of this temporal spectrum presents challenges for studying neopolyploid establishment in natural populations, emphasizing the utility of theory in understanding this process.

An extended period of close interaction between perennial diploids and polyploids may have significant consequences for polyploid evolution during and after establishment. For example, ongoing intercytotype reproductive events could result in selection for assortative mating, driving divergence in floral morphology or phenology between diploids and polyploids (Oswald and Nuismer 2011; Husband et al. 2016). Prolonged intercytotype mating will also promote polyploid establishment when diploid unreduced gamete (UG) production is nonzero, because the probability of two UGs giving rise to a viable polyploid offspring in a diploid-diploid cross is *ug*^2^, but the probability of one UG being involved in a diploid-polyploid cross is *ug* (table S1). Our model is the first to incorporate UG production and perenniality, and we find that a UG production rate of 1-5% is sufficient to ensure polyploid success when diploids and polyploids are perennial, providing polyploids also have a small survival advantage (fig. 2C). This is similar to estimates of UG frequencies in natural populations (0.05-2%; Ramsey and Schemske 1998; Kreiner et al. 2017), and is substantially lower than the UG production rate of ∼17% required for non-selfing annuals with random mating (Felber 1991; Husband 2004), or 10% for annuals with self-fertilization and low inbreeding depression (Rausch and Morgan 2005). UG production may then be less of a limiting step for polyploid formation among perennials where reproductive events are spread over multiple generations.

### CLONALITY INFLUENCES THE ESTABLISHMENT PROCESS

When all else is equal between cytotypes, simply being clonal is not enough to ensure establishment success under any scenario we investigated – some small advantage over diploids is needed (e.g., higher ramet production). In our model, increased clonality confers high establishment success, as in mixed diploid-tetraploid *Chamerion angustifolium* (fireweed), where neotetraploids exhibit immediate increases in clonal rootbud production (Van Drunen and Husband 2018*b*). Conversely, decreased clonal investment in polyploids results in no establishment (fig. 4), a pattern consistent with the lack of natural cytotype variation in *Fragaria vesca* (woodland strawberry), where artificial neotetraploids have lower stolon production (Van Drunen and Husband 2018*a*). These remain the only species in which neopolyploid investment in clonal reproduction has been measured, but if this pattern applies more generally, it would suggest that the production of clonal ramets is an important factor in polyploid establishment (Spoelhof et al. 2017).

Perenniality and clonal reproduction have both been implicated in polyploid success in natural populations (Rice et al. 2019; Van Drunen and Husband 2019), but few studies have contrasted their importance in the early stages of polyploid evolution. Here, we show that polyploids with a combination of perenniality and clonal reproduction generally have greater establishment success than polyploids that are strictly sexual. In some scenarios, perenniality alone was enough to ensure polyploid establishment (e.g., when autogamy was high). While clonal reproduction did not affect final establishment probability in these simulations, it still significantly altered the establishment process. For instance, polyploids were able to spread into the diploid population faster when clonal, even when seedlings and ramets were equal competitors (fig. S10), but this effect was heavily moderated by clonal strategy. Spoelhof et al. (2020) speculated that rapid clonal expansion may increase establishment probability, but result in genetically homogeneous polyploid populations. We confirm this outcome, particularly for clonal strategies that produce many ramets with high lateral spread (fig. 5B). Here, it is possible that polyploids may enjoy short-term success, but may not survive in the long-term if the lack of genetic variability in the polyploid population leads to reduced evolutionary potential (Muller 1932; Pan and Price 2002; Barrett 2015; Herben et al. 2016). According to our results, polyploids conforming to a large-clumped clonal architecture may be the most successful if they spread quickly while maintaining higher genet diversity than polyploids with more dispersible ramets (fig. 5B). Nonetheless, spreading architectures may perform well in environments with high temporal variability or frequent disturbance where slower expansion could result in the stochastic loss of all polyploids (Čertner et al. 2017), especially if genetic uniformity is counteracted by recurrent polyploid formation, immigration, or high rates of vegetative somatic mutation between ramets (Yu et al. 2020).

Our model demonstrates that spatial clonal architecture has a large effect on the polyploid establishment process, but that no single clonal strategy is superior under all circumstances. Similarly, phylogenetic comparative studies have found mixed support for associations between particular clonal strategies and polyploid occurrence, depending on the group of species being studied (Herben et al. 2017; Van Drunen and Husband 2019). Overall, given the wide diversity, evolutionary lability, and multi-functional nature of clonal modes across the angiosperms (Herben and Klimešová 2020), it seems unlikely that there is a one-size-fits-all hypothesis describing the relationship between clonal reproduction and polyploidy. For example, bulbs or corms (generally “clumped” clonal architectures) are involved in resource storage (Klimeš et al. 1997), which may affect over-winter survival or the timing and extent of sexual reproduction in the next growing season. Thus, polyploid establishment may indeed be successful due to clonality in species with these strategies, but for significantly different reasons than explored in our current model.

### CLONAL ARCHITECTURE & MATING

Clonal reproduction may enhance polyploid establishment by creating local polyploid majorities (Baack 2005; Spoelhof et al. 2020), but previous models have overlooked one of the key aspects of this polyploidy-promoting mechanism: geitonogamous self-fertilization. Geitonogamous self-fertilization between different flowers on the same plant, or between clonal ramets, is often regarded as a negative and non-adaptive side-effect of outcrossing (Lloyd 1992; Eckert 2000). Geitonogamy generally involves mating costs, as it requires the same pollination process as would occur with outcrossing, but it reduces outcross siring success and results in pollen (Harder and Barrett 1995; Lau et al. 2008) and seed discounting (Lloyd 1992) when there is any inbreeding depression. But geitonogamy is rarely considered in the complete absence of compatible mates, where self-fertilization may be the only recourse for producing viable sexual offspring. Our model shows that geitonogamous self-fertilization between clonal ramets can confer a significant fitness advantage to establishing polyploids by enabling them to acquire same-cytotype mates, revealing a situation where geitonogamy might be adaptive. Here, when a new polyploid is instantly reproductively isolated from its progenitors, the costs typically associated with pollen and seed discounting via geitonogamy instead apply to intercytotype outcrossing that does not produce viable recruits.

Rates of geitonogamy are expected to vary with clonal architecture (Charpentier 2002; Matsuo et al. 2014; Van Drunen et al. 2015). We can clearly see this in our model, where the long reach of dispersed polyploid genets enables ramet recruitment over a wide area, leading to large genets and high rates of geitonogamy. In contrast, seedlings are equal or better dispersers than ramets when clonal architecture is clumped (fig. S1), resulting in smaller genets, more sexual recruits, and more frequent intracytotype outcrossing than geitonogamous self-fertilization (fig. 6). These mating differences contribute to establishment speed and overall establishment probability, especially when pollen and seed dispersal distances are long and sexual reproduction is not dominated by autogamy. Across all clonal strategies, however, geitonogamy substantially reduced ovule waste to intercytotype mating early in the establishment process (fig. 6).

We might then predict that neopolyploids will experience strong selection on traits that increase geitonogamy, such as ramet production (Hu et al. 2015), clonal architecture (Van Drunen and Husband 2018*a*), or floral display size (Vamosi et al. 2007). But as polyploids spread throughout the population and intracytotype outcrossing becomes more probably, selection may reverse and instead favour outcross sexual reproduction. There is some evidence for this occurring in autotetraploid *Chamerion angustifolium*, where initial WGD-driven increases in synthetic neotetraploid rootbud production is significantly reduced in naturally-occurring tetraploids (Van Drunen and Husband 2018*b*; Walczyk and Hersch-Green 2019). In our model we saw a shift towards recruiting outcrossed seedlings as polyploids became common (fig. 6), but we could not evaluate selection on reproductive strategy directly because we did not include phenotypic trait evolution. Future work that allows for heritable variation in reproductive strategy and sex allocation, and resource-based trade-offs between reproductive modes could therefore be particularly insightful.

Clonal reproduction and geitonogamous self-fertilization may be less likely to facilitate polyploid success when pollen is limited and pollen discounting is high (Yamauchi 2006), or when ramet recruitment into the population is low. In our model, we assumed that clonal ramets were better competitors than seedlings, and so polyploid ramets were recruited much more often than sexual offspring, even when they were less common in the recruitment pool (fig. 6). In contrast, when ramets and seedlings were equal competitors in the exponential ramet dispersal scenarios there were no differences in establishment probability between clonal and nonclonal polyploids, though fixation times were still substantially shorter (figs. S8, S10). Ramet competitive superiority is realistic for most clonal species, where ramet recruits tend to be more common than seedlings within populations (Eriksson 1993; Vallejo-Marín et al. 2010; Vandepitte et al. 2010; Johnson et al. 2020), but the validity of this assumption may vary for different taxa, and ecological or demographic conditions. For instance, considering only within-population processes ignores the role of seedlings in long-distance colonisation events and meta-populations dynamics (Winkler and Fischer 2001; Griswold 2021; but see Scherrer et al. 2017), and thus polyploids arising from diploids that occur in patchy habitats might benefit from more dispersible sexual offspring than locally-restricted clonal propagules.

### CONCLUSIONS

Whether perenniality and clonal reproduction promote polyploid establishment depends on multiple factors, including the phenotypic effects of WGD, local habitat availability, clonal strategy, mating patterns and pollination system, inbreeding depression, and more. Ultimately, determining whether the processes in our model occur in natural populations will require fine-scale genetic data and extensive sampling, and may not be possible in polyploid systems that are long established. Further models exploring a variety of situations (e.g., clonal vs. sexual reproduction and polyploid establishment in patchy habitats), and experimental studies involving synthetic neopolyploids or very young natural neopolyploids (e.g., tetraploid Mimulus guttatus in the Shetland Isles; Simón-Porcar et al. 2017) will be key to unraveling the individual and interactive roles of perenniality and clonality in early polyploid evolution.

## ACKNOWLEDGEMENTS

We thank B.C. Husband for input on study concept. Funding for this study was provided by the Natural Sciences and Engineering Research Council (NSERC) through a Postdoctoral Fellowship to W.E.V., and a Discovery Grant to J.F.

## DATA AND CODE AVAILABILITY

All of the code needed to run the simulation model in this article is available online at: https://github.com/wevandrunen/autopolyploid-establishment-lifehistory-clonalarchitecture.

## AUTHOR CONTRIBUTION STATEMENT

W.E.V. and J.F. conceived of the study. W.E.V. developed the model, and analyzed the output. W.E.V. wrote the manuscript with input from J.F.

## Notes

https://github.com/wevandrunen/autopolyploid-establishment-lifehistory-clonalarchitecture

### Competing Interest Statement

The authors have declared no competing interest.

