## Supplemental Materials for "Autopolyploid establishment depends on life history strategy and the mating outcomes of clonal architecture"

### List of Tables:


| | |
|---|---|
| **Table S1** | Seeds produced in a population containing both diploid and tetraploid individuals categorized into 22 cross-types, dependent on the identity of pollen and ovule sources for a given mating event. |


### List of Figures:


| | |
|---|---|
| **Figure S1** | Pollen, seed, and ramet dispersal density functions. |
| **Figure S2** | Mean time to tetraploid fixation or exclusion for a range of diploid and tetraploid survival probabilities. |
| **Figure S3** | Tetraploid establishment patterns in a population with reduced density. |
| **Figure S4** | Tetraploid establishment patterns when selfed-seed inviability is unequal between cytotypes. |
| **Figure S5** | Tetraploid establishment patterns when unreduced gamete (UG) production is non-zero. |
| **Figure S6** | Tetraploid frequency and mixed-cytotype population stability after 500 generations under non-zero UG production. |
| **Figure S7** | Tetraploid establishment patterns for different clonal strategies, across the core dispersal-inviability parameter set. |
| **Figure S8** | The effects of reduced tetraploid selfed-seed inviability on tetraploid establishment between nonclonal, normal ramet dispersal, and exponential ramet dispersal scenarios for equal life history strategies. |
| **Figure S9** | The influence of interactions between life history and clonal strategies on tetraploid establishment patterns. |
| **Figure S10** | The influence of interactions between life history and clonal strategies on tetraploid establishment patterns, when ramet dispersal is exponential. |
| **Figure S11** | Tetraploid establishment probability when tetraploids and diploids have different clonal architectures. |
| **Figure S12** | Tetraploid mating outcomes and offspring recruitment when ramet dispersal is exponential. |

**Table S1**: Seeds produced in a population containing both diploid and tetraploid individuals can be categorized into 22 cross-types, dependent on the identity of pollen and ovule sources for a given mating event. Self-fertilized seeds are inviable with probability $k_2$ (diploids) or $k_4$ (tetraploids), and gametes are unreduced with probability *ug*. Triploid offspring are not viable. See Figure 1 and the main text for parameter descriptions.

| Cross-type | | | Ovule | Pollen | Probability | | Offspring ploidy | Viable |
|---|---|---|---|---|---|---|---|---|
| | | | | | $ug = 0$ | $ug \neq 0$ | | |
| *Ovule Source is Diploid:* | | | | | | | | |
| Self-fertilization (within shoot) | 1 | Inviable self | 1n | 1n | $k_2$ | $(1-ug)^2 k_2$ | 2x | |
| | 2 | Viable self | 1n | 1n | $1-k_2$ | $(1-ug)^2(1-k_2)$ | 2x | ✓ |
| | 3 | Single UG self | 1n / 2n | 2n / 1n | 0 | $2(ug)(1-ug)$ | 3x | |
| | 4 | Inviable double UG self | 2n | 2n | 0 | $ug^2 k_2$ | 4x | |
| | 5 | Viable double UG self | 2n | 2n | 0 | $ug^2(1-k_2)$ | 4x | ✓ |
| Self-fertilization (between ramet) | 6 | Inviable self | 1n | 1n | $k_2$ | $(1-ug)^2 k_2$ | 2x | |
| | 7 | Viable self | 1n | 1n | $1-k_2$ | $(1-ug)^2(1-k_2)$ | 2x | ✓ |
| | 8 | Single UG self | 1n / 2n | 2n / 1n | 0 | $2(ug)(1-ug)$ | 3x | |
| | 9 | Inviable double UG self | 2n | 2n | 0 | $ug^2 k_2$ | 4x | |
| | 10 | Viable double UG self | 2n | 2n | 0 | $ug^2(1-k_2)$ | 4x | ✓ |
| Intercytotype | 11 | Intercytotype outcross | 1n | 2n | 1 | $1-ug$ | 3x | |
| | 12 | Single UG intercytotype outcross | 2n | 2n | 0 | $ug$ | 4x | ✓ |
| Intracytotype | 13 | Intracytotype outcross | 1n | 1n | 1 | $(1-ug)^2$ | 2x | ✓ |
| | 14 | Single UG intracytotype outcross | 1n / 2n | 2n / 1n | 0 | $2(ug)(1-ug)$ | 3x | |
| | 15 | Double UG intracytotype outcross | 2n | 2n | 0 | $ug^2$ | 4x | ✓ |
| *Ovule Source is Tetraploid:* | | | | | | | | |
| Self-fertilization (within shoot) | 16 | Inviable self | 2n | 2n | $k_4$ | $k_4$ | 4x | |
| | 17 | Viable self | 2n | 2n | $1-k_4$ | $1-k_4$ | 4x | ✓ |
| Self-fertilization (between ramet) | 18 | Inviable self | 2n | 2n | $k_4$ | $k_4$ | 4x | |
| | 19 | Viable self | 2n | 2n | $1-k_4$ | $1-k_4$ | 4x | ✓ |
| Intercytotype | 20 | Intercytotype outcross | 2n | 1n | 1 | $1-ug$ | 3x | |
| | 21 | Single UG intercytotype outcross | 2n | 2n | 0 | $ug$ | 4x | ✓ |
| Intracytotype | 22 | Intracytotype outcross | 2n | 2n | 1 | 1 | 4x | ✓ |

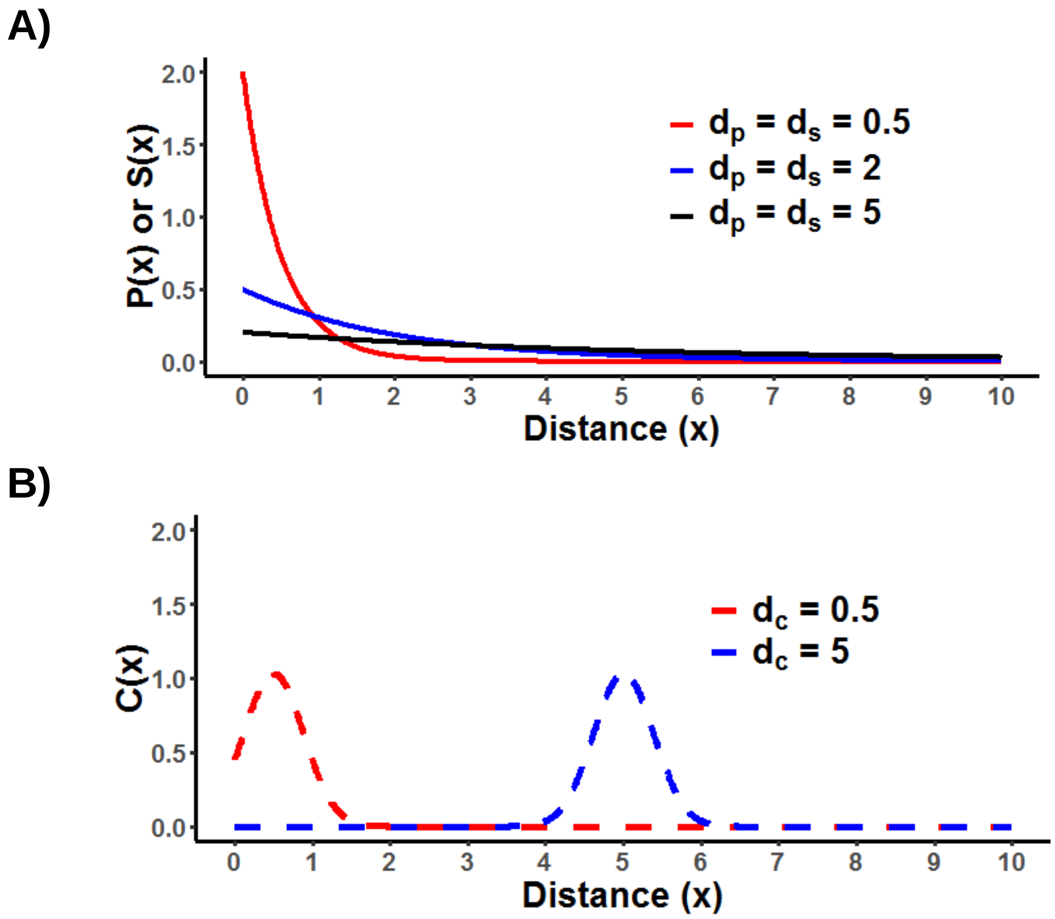


**Figure S1**: Probability density functions for pollen, seed, and ramet dispersal. A) Exponential probability density functions for pollen *P(x)* and seed *S(x)* dispersal (Eq. 1 in the main text), for short ($d_p = d_s = 0.5$), mid ($d_p = d_s = 2$), and far ($d_p = d_s = 5$) average dispersal distances. B) Gaussian (normal) probability function for clonal ramet *C(x)* dispersal (Eq. 2 in the main text), for short ($d_c = 0.5$, *Phalanx*) and spreading ($d_c = 5$, *Guerrilla*) clonal architectures

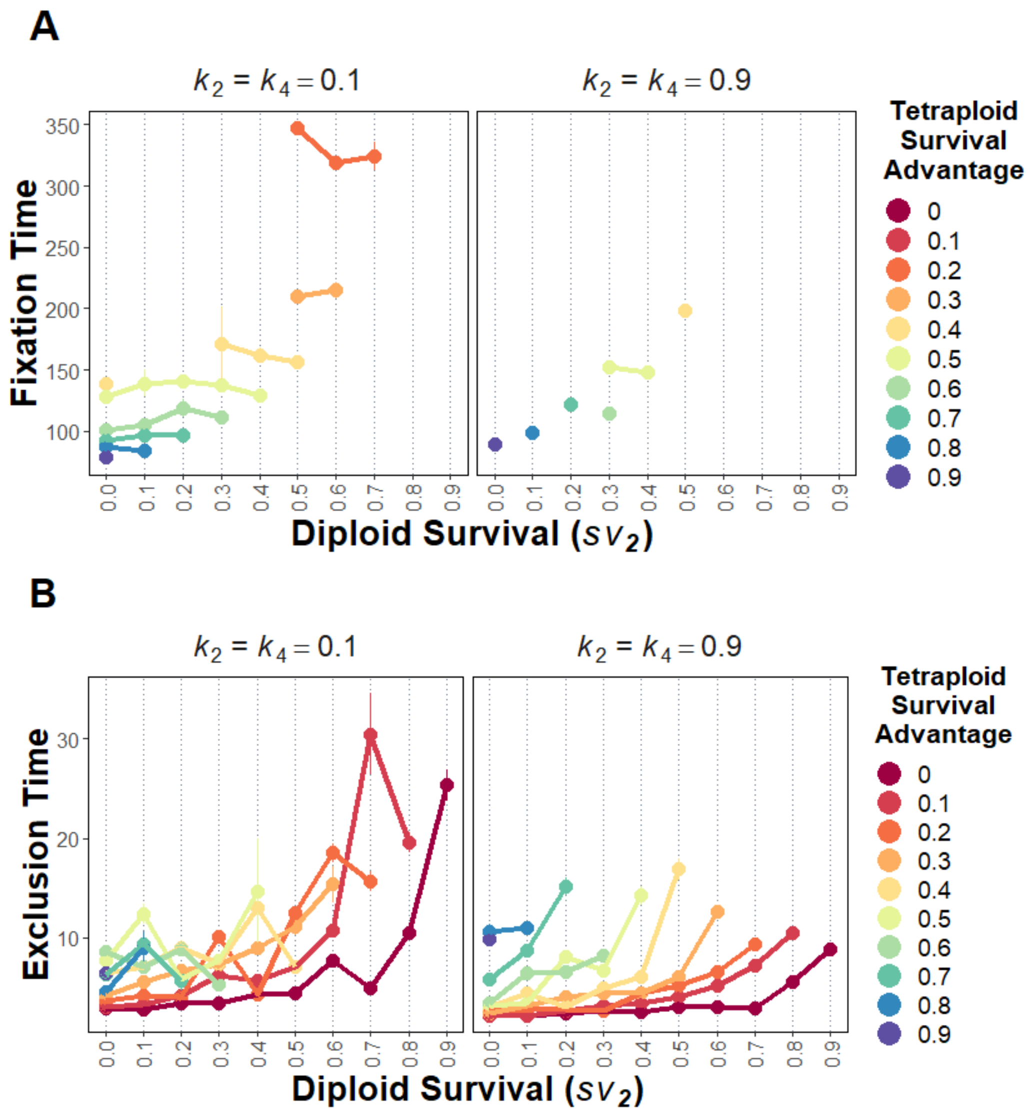


**Figure S2**: The effect of life history strategy on mean time (± SE) to tetraploid fixation (A) or exclusion (B), for a range of diploid and tetraploid per generation survival probabilities ($sv_2$, $sv_4$). Labels above plots indicate selfed-seed inviability ($k = k_2 = k_4$). For all models $d_p = d_s = 0.5$, and $ug = 0$.

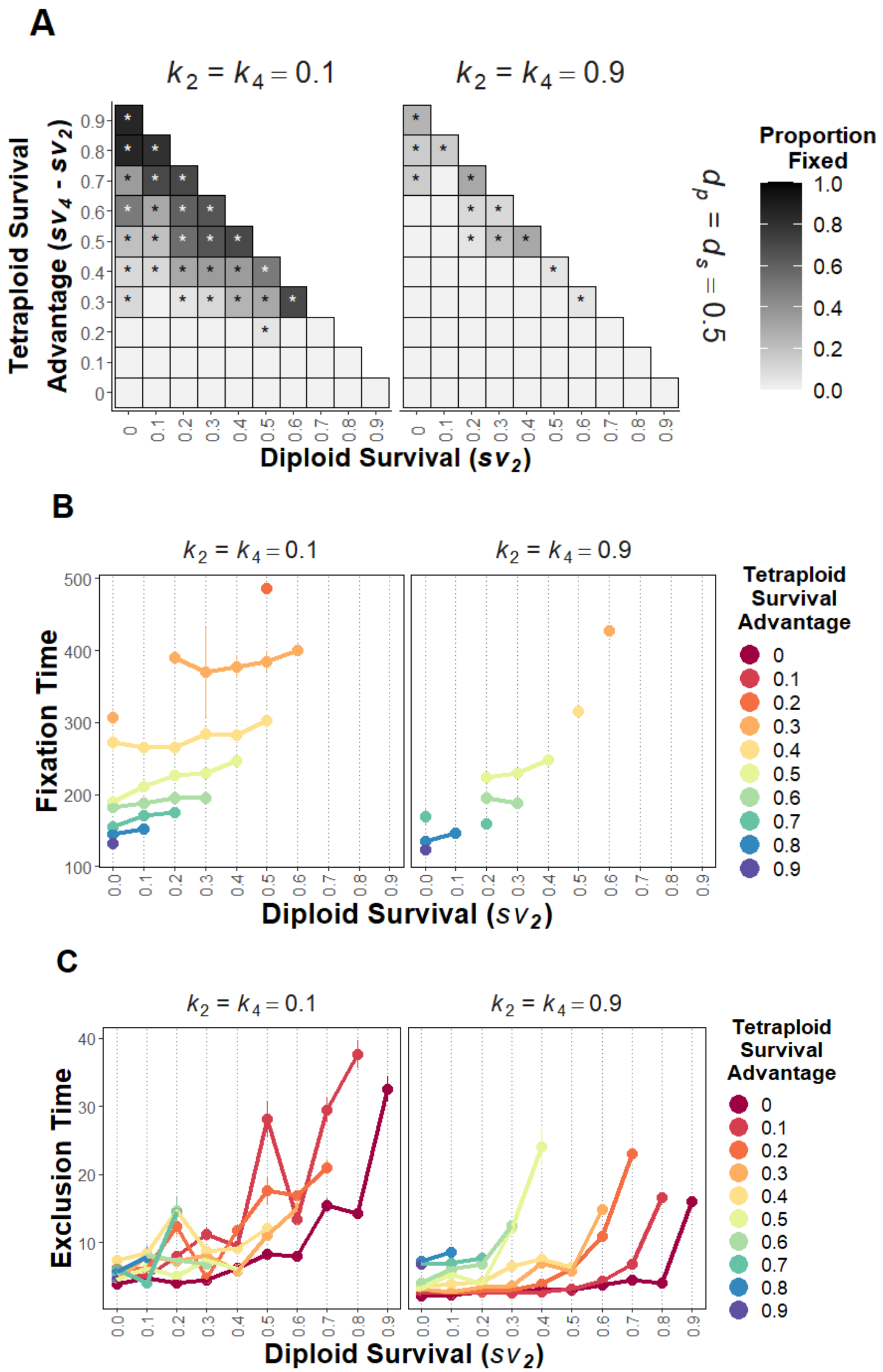


**Figure S3**: The effect of life history strategy on polyploid establishment dynamics on a larger population lattice with lower population density ($D$ = 40, $N$ = 900), for a range of diploid and tetraploid per generation survival probabilities ($sv_2$, $sv_4$). A) The proportion of simulation runs where tetraploids go to fixation. Asterisks denote parameter combinations where at least one simulation replicate resulted in tetraploid fixation. B) Mean number of generations to tetraploid fixation (± SE) across simulation replicates. C) Mean time to tetraploid exclusion (± SE). Labels above plots indicate selfed-seed inviability ($k = k_2 = k_4$). For all models $d_p = d_s = 0.5$, and $ug = 0$.

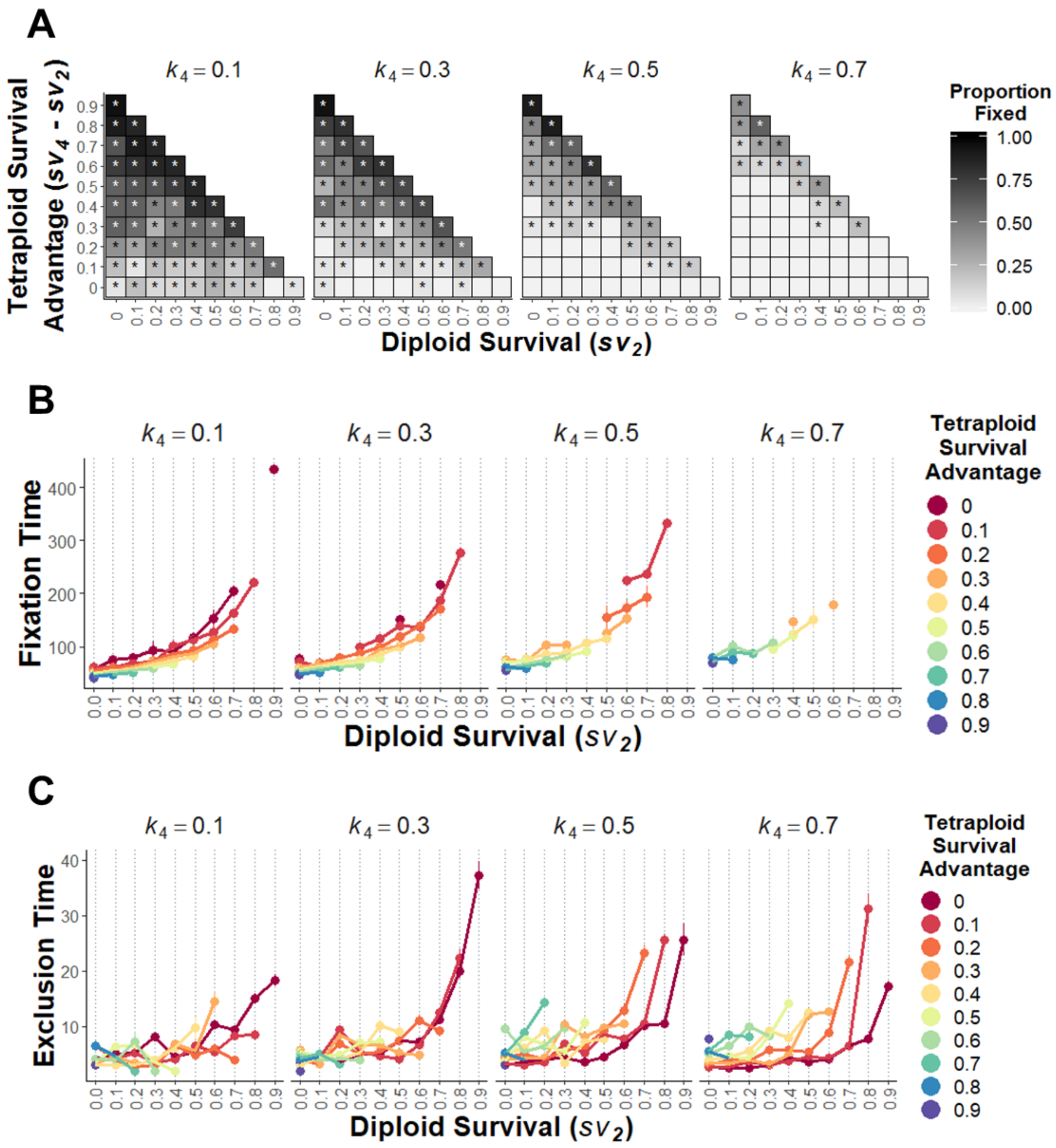


**Figure S4:** The effect reduced tetraploid selfed seed inviabilty ($k_4$) relative to diploid selfed seed inviability ($k_2$) on tetraploid establishment, are for a range of diploid and tetraploid per generation survival probabilities ($sv_2$, $sv_4$). Labels above panels indicate values of $k_4$, and $k_2$ = 0.9 for all scenarios. A) The proportion of simulation runs where tetraploids go to fixation. Asterisks denote parameter combinations where at least one simulation replicate resulted in tetraploid fixation. B) Mean number of generations to tetraploid fixation (± SE) across simulation replicates. C) Mean time to tetraploid exclusion (± SE). For all models $d_p$ = $d_s$ = 0.5, and $ug$ = 0.

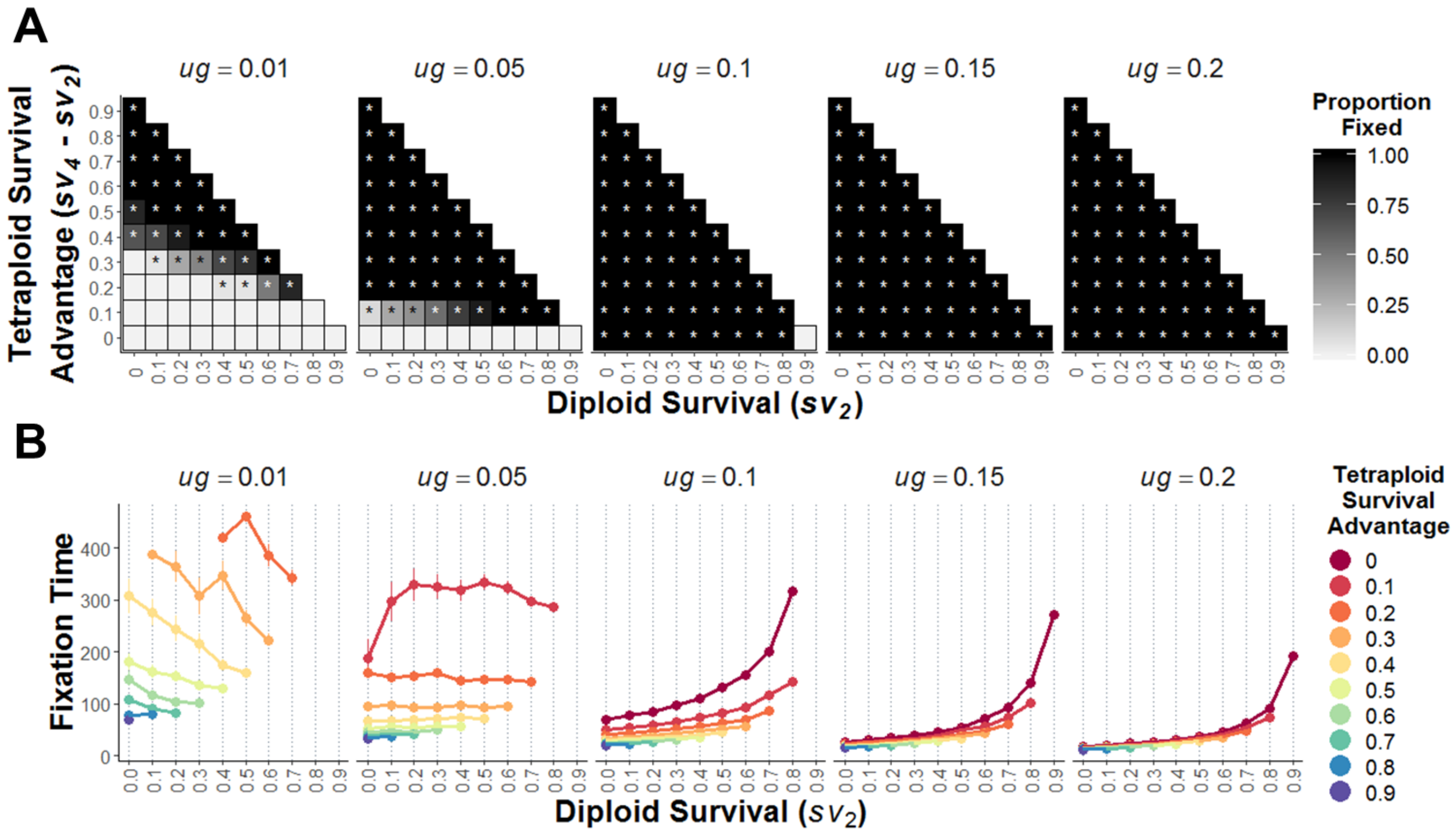


**Figure S5:** Polyploid establishment under non-zero unreduced gamete (UG) production, for a range of diploid and tetraploid per generation survival probabilities ($sv_2$, $sv_4$). Labels above panels indicate *ug* value. A) The proportion of simulation runs where tetraploids go to fixation. Asterisks denote parameter combinations where at least one simulation replicate resulted in tetraploid fixation. B) Mean time to tetraploid fixation (± SE). Note that tetraploid cannot be excluded from a population when $ug > 0$. For all models $d_p = d_s = 0.5$, and $k_2 = k_4 = 0.1$

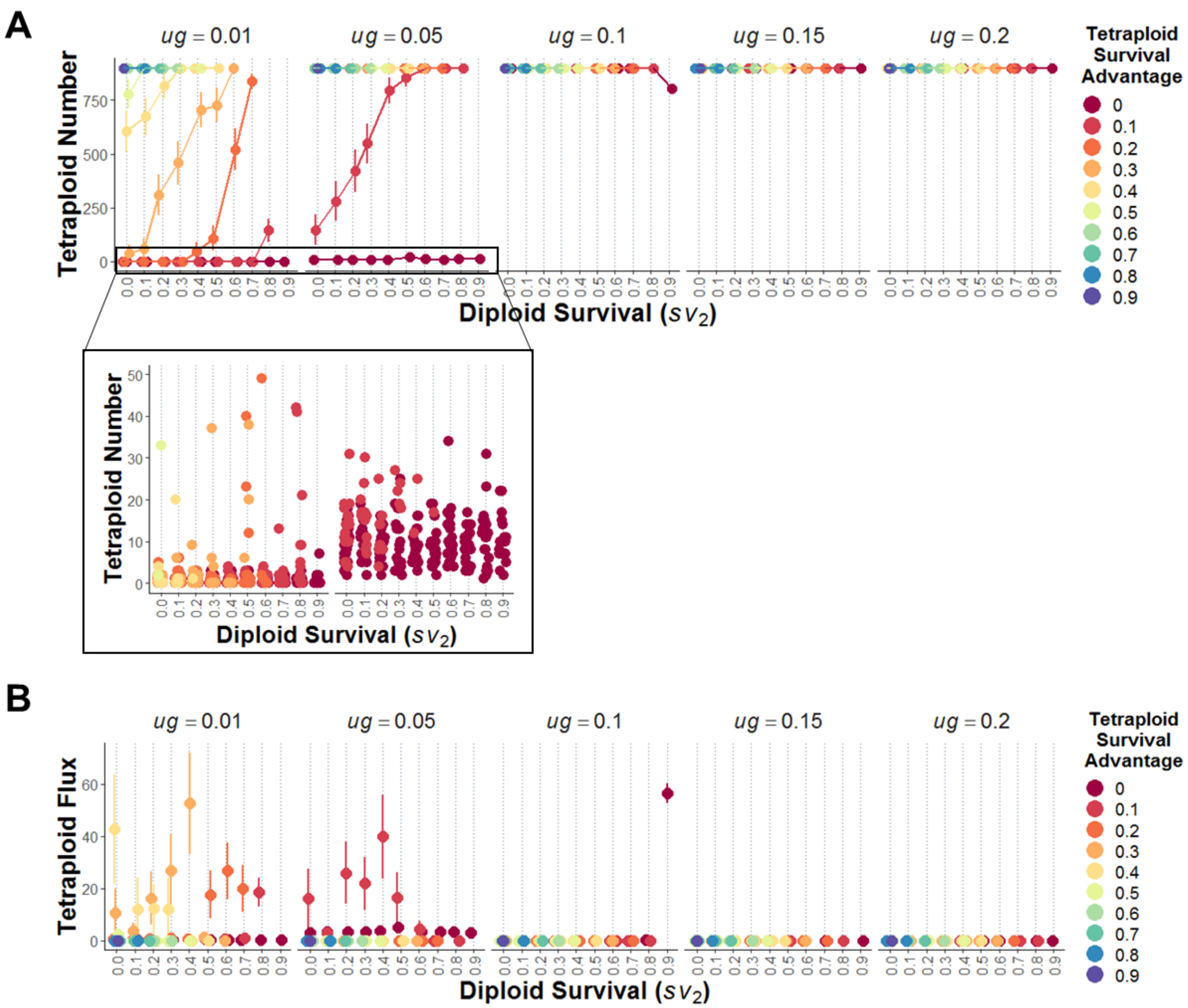


**Figure S6**: Tetraploid population after 500 generations when unreduced gamete production (*ug*) is non-zero, for a range of diploid and tetraploid per generation survival probabilities ($sv_2$, $sv_4$). Labels above panels indicate *ug* value. A) Mean number of tetraploids (± SE) at 500 generations across 20 simulation replicates. Zoomed inset shows number of tetraploids at the end of each replicate. B) Mean fluctuation (± SE) in tetraploid number over the last 100 generations, calculated as the deviation from the mean tetraploid number per generation across replicates. Fluctuations of < 10 individuals per generation are considered to be stable equilibria. For all models $d_p = d_s = 0.5$, and $k_2 = k_4 = 0.1$. Points are slightly horizontally jittered.

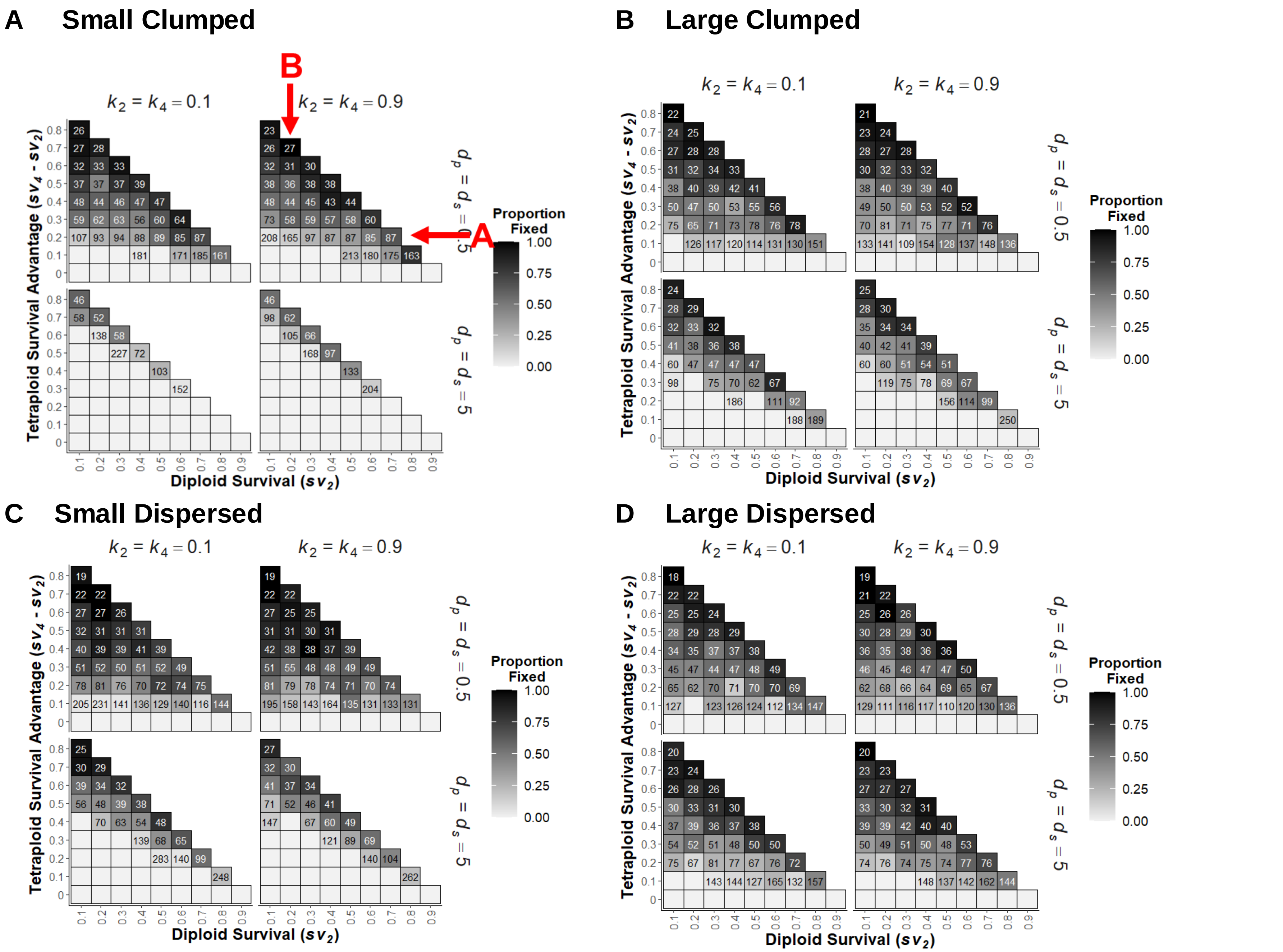


**Figure S7:** The influence of clonal strategy on tetraploid establishment probability across the core dispersal-inviability parameter set, for a range of diploid and tetraploid per generation survival probabilities ($sv_2$, $sv_4$). Clonal strategy is equal between cytotypes. Tile numbers represent average times to fixation. For all models $ug = 0$. Arrows in (A) demonstrate the columns or rows shown in Figure 3A and B.

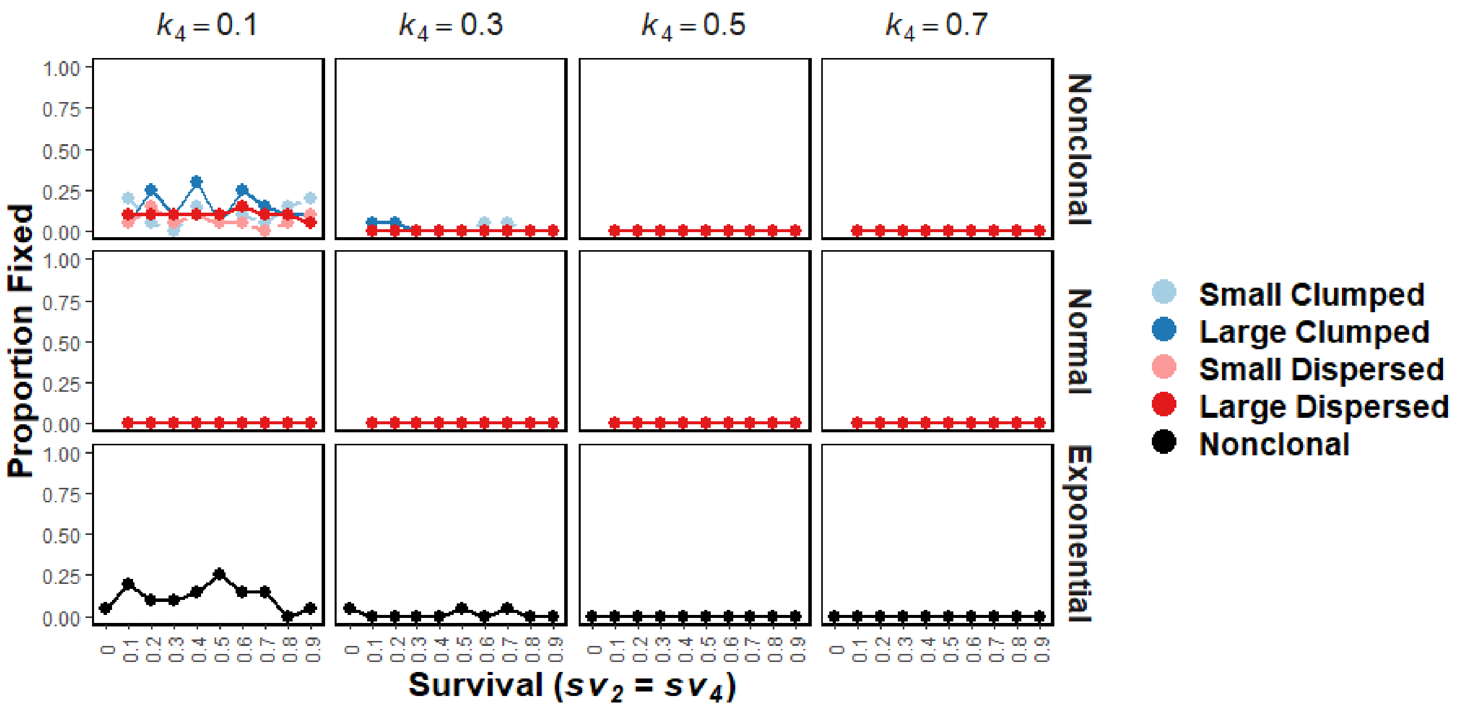


**Figure S8:** Tetraploid establishment probability between the non-clonal scenario, and when ramet dispersal follows either a normal (Gaussian, Figure S1B, Eqn. 2) or exponential.ramet dispersal (Figure S1A, Eqn. 1). Clonal and life history strategies between diploids and tetraploids are equal, but selfed seed inviability for tetraploids is lower than for diploids ($k_2$ = 0.9, $k_4$ varies between columns). For all models $d_p = d_s = 0.5$, and $ug = 0$.

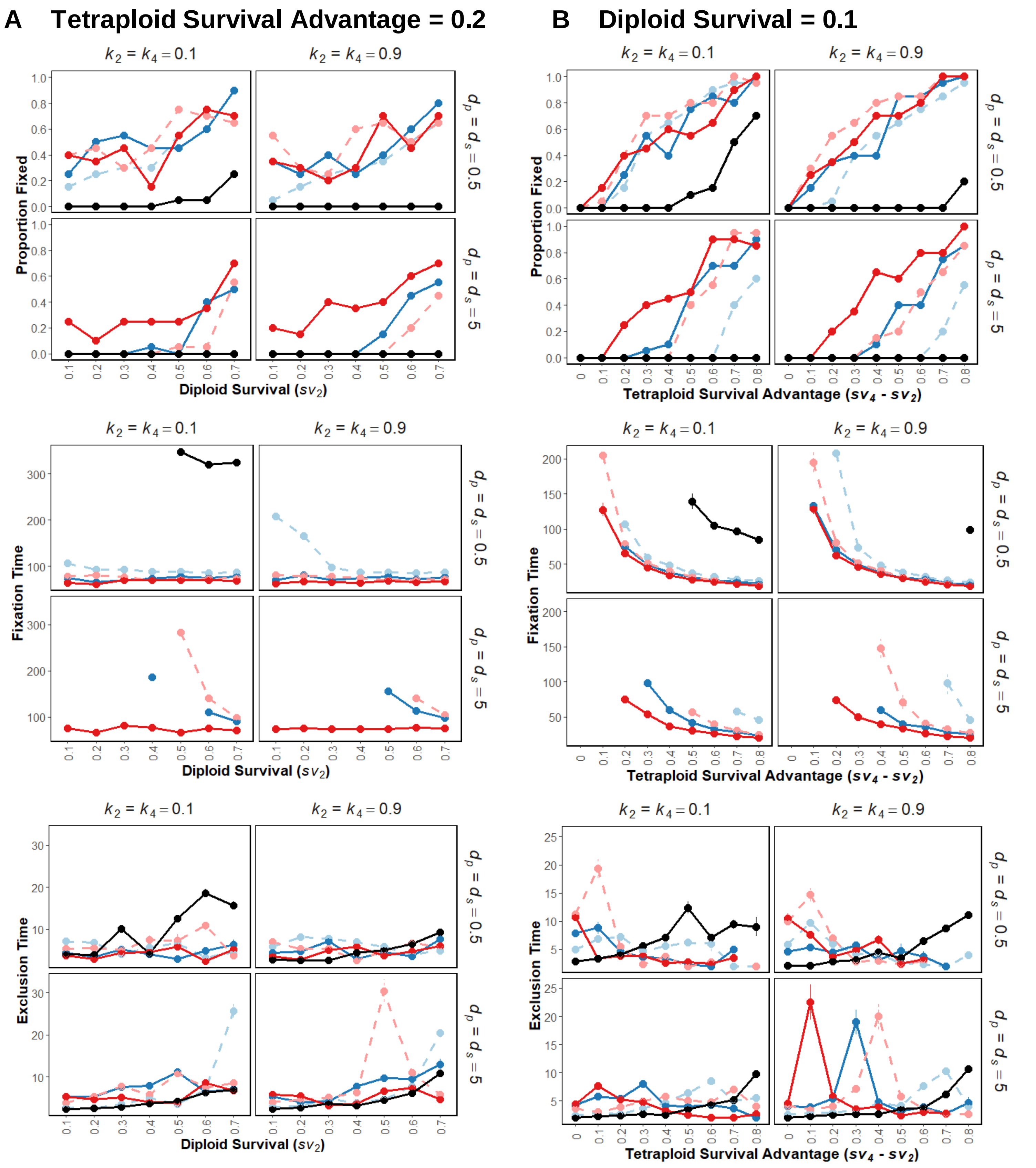


**Figure S9:** Establishment probability, and mean time to tetraploid fixation or exclusion across the core dispersal-inviability parameter set, for scenarios where tetraploid survival advantage is either (A) 0.2 higher than diploid survival (left column), or (B) diploid survival is constant at 0.1 and tetraploid survival can take higher values (right column). Clonal strategy is equal between cytotypes (black = nonclonal, light blue= small clumped, dark blue = large clumped, light red = small dispersed, dark red = large dispersed). For all models *ug* = 0.

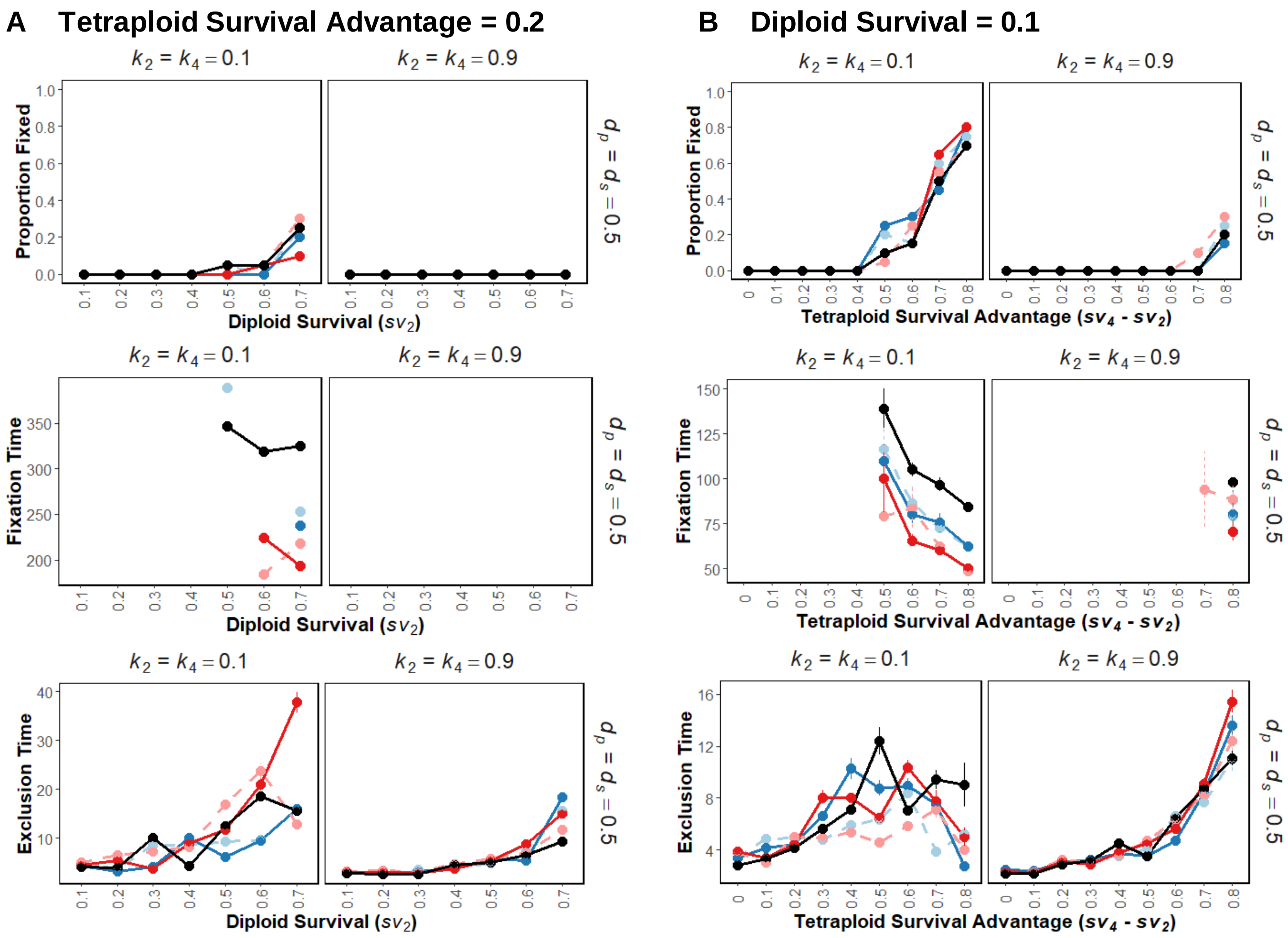


**Figure S10:** Tetraploid establishment, mean time to fixation, and mean time to exclusion under exponential ramet dispersal, for scenarios where tetraploid survival advantage is either (A) 0.2 higher than diploid survival (left column), or (B) diploid survival is constant at 0.1 and tetraploid survival can take higher values (right column). Clonal strategy is equal between cytotypes (black = nonclonal, light blue= small clumped, dark blue = large clumped, light red = small dispersed, dark red = large dispersed). Selfed-seed inviability ($k = k_2 = k_4$) varies along columns. No tetraploid establishment occurred when $d_p = d_s = 2$ or 5. For all models $ug = 0$.

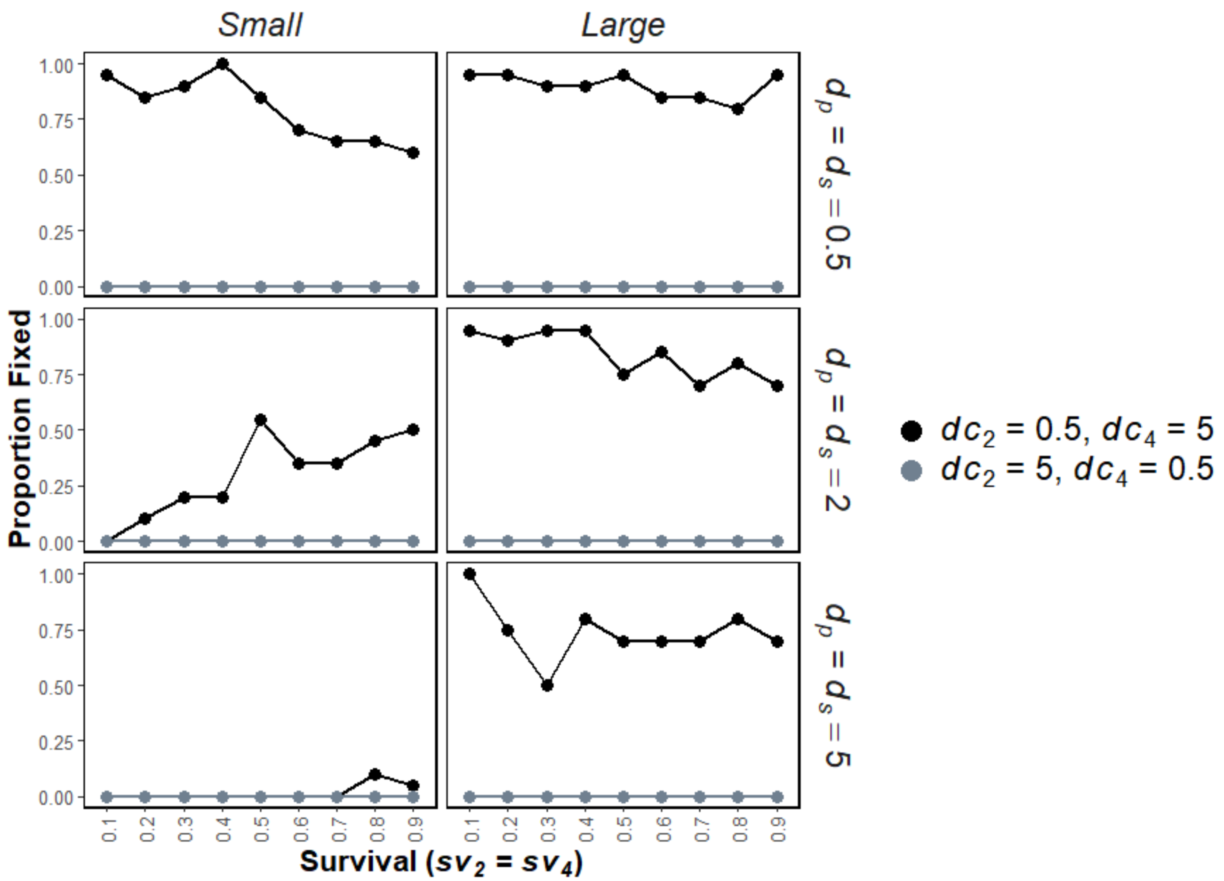


**Figure S11:** Tetraploid establishment probability when cytotypes have different clonal architectures ($d_{C2} \neq d_{C4}$), which may be clumped ($d_{C2}$ = 0.5) or dispersed ($d_{C2}$ = 5). Ramet production is equal between cytotypes, resulting in genets that are small ($c_2 = c_4 = 1$) or large ($c_2 = c_4 = 5$). Life history strategies are equal between cytotypes ($sv_2 = sv_4$), as shown on the x-axis. For all models, selfed-seed inviability is equal between cytotypes ($k_2 = k_4 =$ 0.1), and there is no unreduced gamete production ($ug$ = 0).

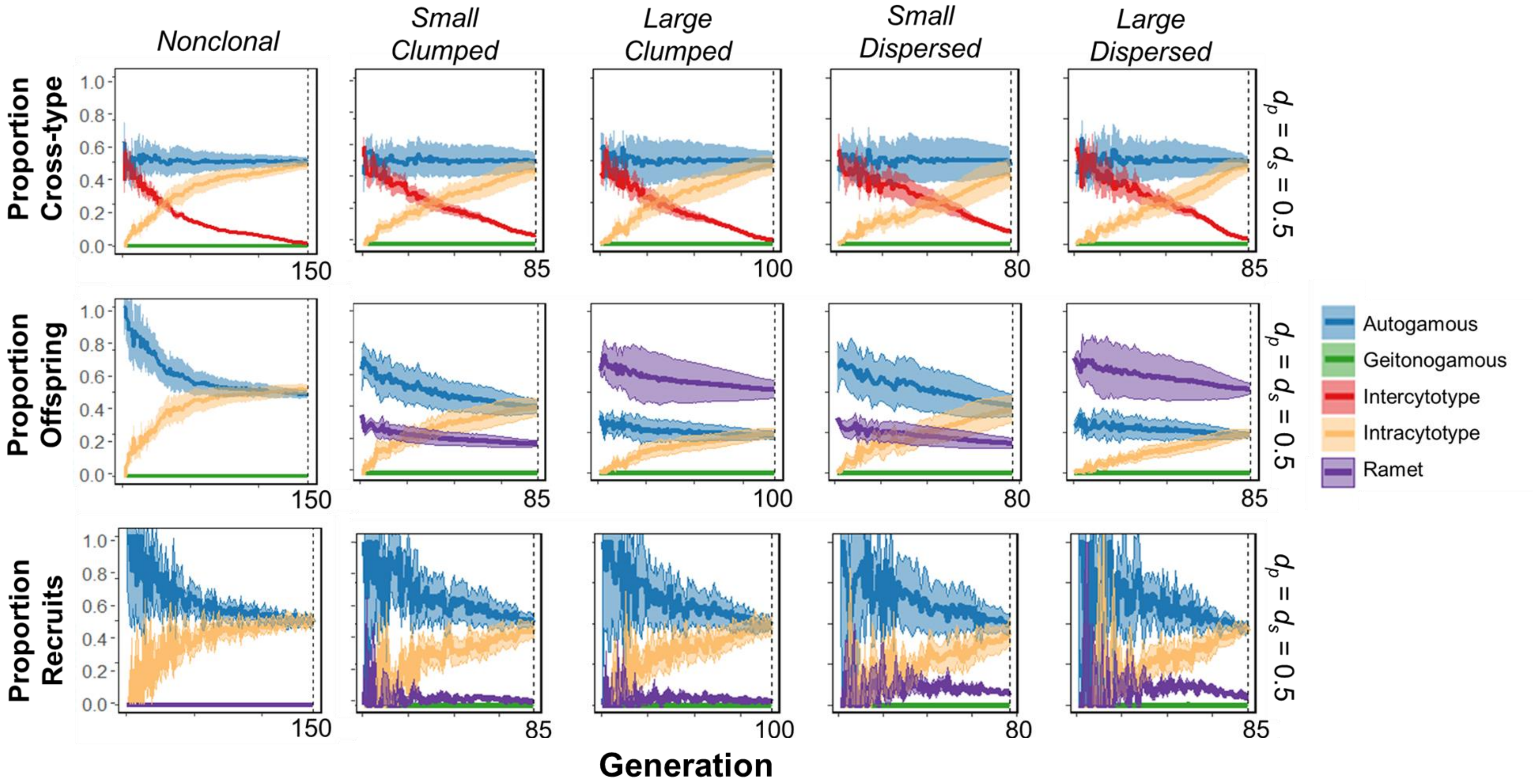


**Figure S12:** Differences in cross-type production, viable offspring production, and offspring recruitment between clonal architectures when ramet dispersal follows an exponential distribution. Clonal strategy varies across columns, and cytotypes have identical architecture (*Clumped* $d_{C2} = d_{C4} = 0.5$; *Dispersed* $d_{C2} = d_{C4} = 5$), and equal ramet production (*Small* $c_2 = c_4 = 1$; *Large* $c_2 = c_4 = 5$). For all models, $sv_2 = 0.5$ and $sv_4 = 0.9$, $d_p = d_s = 0.5$, $k_2 = k_4 = 0.1$, and $ug = 0$. Generations are truncated at the fastest tetraploid fixation time per parameter combination, indicated at the bottom right of each panel.
